# Pulmonary osteoclast-like cells in silica induced pulmonary fibrosis

**DOI:** 10.1101/2023.02.17.528996

**Authors:** Yoshihiro Hasegawa, Jennifer M. Franks, Yusuke Tanaka, Yasuaki Uehara, David F. Read, Claire Williams, Sanjay Srivatsan, Lori B. Pitstick, Nikolaos M. Nikolaidis, Ciara M. Shaver, Huixing Wu, Jason C. Gardner, Andrew R. Osterburg, Jane J. Yu, Elizabeth J. Kopras, Steven L. Teitelbaum, Kathryn A. Wikenheiser-Brokamp, Cole Trapnell, Francis X. McCormack

## Abstract

The pathophysiology of silicosis is poorly understood, limiting development of therapies for those who have been exposed to the respirable particle. We explored the mechanisms of silica-induced pulmonary fibrosis in a mouse model using multiple modalities including whole-lung single-nucleus RNA sequencing. These analyses revealed that in addition to pulmonary inflammation and fibrosis, intratracheal silica challenge induced osteoclast-like differentiation of alveolar macrophages and recruited monocytes, driven by induction of the osteoclastogenic cytokine, receptor activator of nuclear factor-κB ligand (RANKL) in pulmonary lymphocytes and alveolar type II cells. Furthermore, anti-RANKL monoclonal antibody treatment suppressed silica-induced osteoclast-like differentiation in the lung and attenuated silica-induced pulmonary fibrosis. We conclude that silica induces osteoclast-like differentiation of distinct recruited and tissue resident monocyte populations, leading to progressive lung injury, likely due to sustained elaboration of bone resorbing proteases and hydrochloric acid. Interrupting osteoclast-like differentiation may therefore constitute a promising avenue for moderating lung damage in silicosis.

**One Sentence Summary:** Silica induces the alveolar epithelium to reprogram recruited and resident pulmonary myeloid cells to become osteoclasts that contribute to pulmonary fibrosis.

## INTRODUCTION

Crystalline silica is the most ubiquitous mineral on earth (*1-4*). The inhalation of silica particles causes silicosis, a chronic interstitial lung disease that results in progressive pulmonary dysfunction (*1-4*). As one of the most important occupational diseases in both developing and developed nations, silicosis is estimated to impact 2.2 million workers in USA (*5*), 2 million workers in European Union (*6*), and more than 23 million workers in China (*4, 7, 8*).

Pathological subtypes of fibrosis in silicosis include simple nodular silicosis, progressive massive fibrosis, and diffuse interstitial fibrosis (*2, 4*). Silica inhalation is also associated with several life-threatening comorbidities, such as acute silicoproteinosis, pulmonary tuberculosis, chronic obstructive pulmonary disease (COPD), and lung cancer (*1, 4, 9, 10*). The pathogenesis of silicosis is complex but clearly involves engagement of macrophage scavenger receptors and activation of the inflammasome, release of reactive oxygen species, matrix remodeling enzymes, and proinflammatory and profibrotic cytokines and chemokines (*4, 11, 12*).

The advent of single nucleus RNA sequencing and the availability of powerful resources such as the Human Lung Atlas and LungMap provide an opportunity to dissect complex disease mechanisms at levels of resolution that have not previously been possible. Here, we employ single nucleus RNA sequencing in combination with multiple experimental methods to perform a longitudinal study of silica exposure in whole mouse lung. We report evidence that particulate induced osteoclastic differentiation of monocytes and macrophages plays a key role in silica induced pulmonary fibrosis.

Osteoclasts are multinucleated giant cells (MNGC) that mediate bone resorption and maintain normal bone health. MNGC originating from hematopoietic, myeloid, multipotential precursors differentiate into mature osteoclasts under the influence of two cytokines: macrophage colony stimulating factor (M-CSF; CSF1) and receptor activator of nuclear factor-κB ligand (RANKL; TNFSF11) (*13-16*). Osteoclast formation and function can also be stimulated by inflammatory cytokines, tumor necrosis factor alpha (TNF-α), interleukin (IL)-1β, and IL-6 (*17, 18*). Osteoclasts attach to bone, assemble cytoskeletal actin rings and seal off an area through binding of integrin β3 to osteopontin (SPP1) and other bone ligands. Active secretion of H^+^ and Cl^-^ by ATPase H+ Transporting v0 Subunit d2 (ATP6V0D2) and chloride voltage-gated channel 7 (ClCN7) degrade the mineral components of bone and expose the bone matrix to Cathepsin K (CTSK) and other secreted proteases (*19-21*). Although occasionally found outside of bone, osteoclasts have not been previously reported to have physiologic roles in the lung or pathophysiologic roles in pulmonary fibrosis.

## RESULTS

### Pulmonary inflammation and fibrosis in human and murine silicosis

Inhalation of silica particles is known to result in pulmonary inflammation and fibrosis (Fig. 1). Lung specimens obtained from patients with varying degrees of silicosis related pathology were analyzed. Lung tissues from Patient 1 had histopathologic features of late-stage lung disease with extensive fibrosis, collagenized silicotic nodules and patchy chronic inflammation. In contrast, lung tissues from Patient 2 showed less extensive fibrosis with prominent dust macules representing a common reaction to inhaled dust along with rare, small collagenized nodules.

**Fig. 1:**
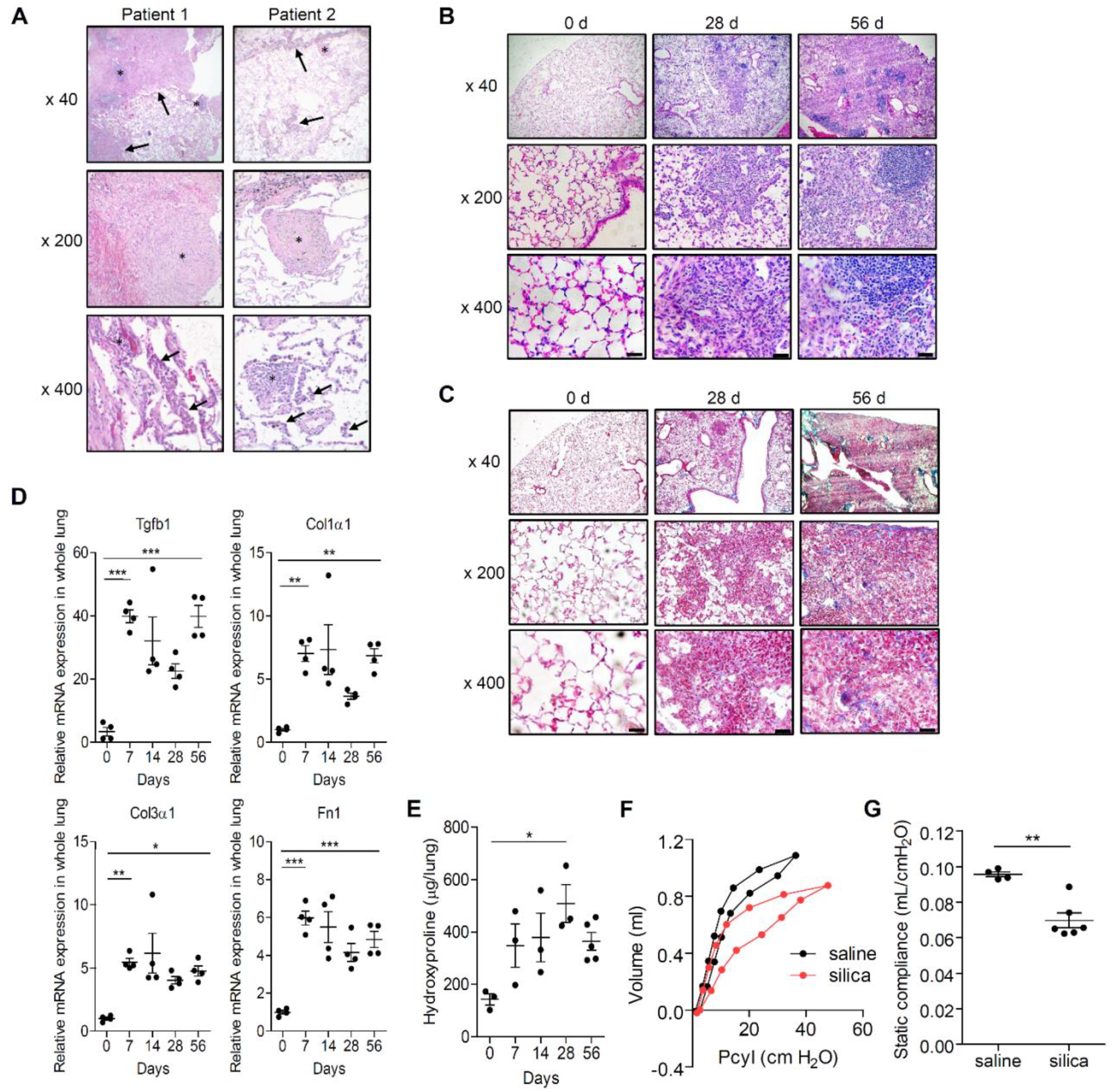
Silicosis features of progressive fibrosis, silicotic nodules, and infiltrating monocytes are recapitulated in the mouse model. Surgical lung specimens from patients with silicosis were analyzed by H & E staining (A). Lung sections from Patient 1 had extensive fibrosis (arrows, top panel) with lymphocyte predominant inflammation (*, top panel), collagenized silicotic nodules (*, middle panel), and surrounding alveoli with septal inflammation (*, bottom panel) and intraalveolar macrophage accumulations (arrows, bottom panel). Lung sections from Patient 2 had focal fibrosis with prominent dust macules and accumulations of pigment containing macrophages (arrows, top panel), rare small collagenized silicotic nodules (*, top and middle panel), and alveoli with lymphocyte predominant septal inflammation (*, bottom panel) and intra-alveolar macrophage aggregates (arrows, bottom panel). Silica particles (2.5 mg) were administered to C57BL/6J mice by the intratracheal route (i.t.). Mice were sacrificed at the indicated times in days after challenge. Paraffin-embedded lung sections were stained with H & E (B) and Masson’s trichrome reagent (C) (Scale bar, 20 μm). (D) Whole lung homogenates were collected from mice at the indicated times in days after silica i.t. administration, and fibrosis-related gene expression was assessed by RT-qPCR for Tgfb1, Col1α1, Col3α1, Fn1 (N = 4 mice per group). (E) Hydroxyproline levels in whole lungs were quantified (N = 3-5 mice per group). (F) At 8 weeks post silica challenge (5 mg, i.t.), pulmonary function tests were performed. The representative curves of pressure in the cylinder (Pcyl)/volume from the two groups and (G) compliance measurements were made using the forced oscillation method and plotted as described in Materials and Methods (N=4-6 mice per group). \**P<*0.05, \*\**P<*0.01 and \*\*\**P<*0.001.

As a model of silicosis, we challenged mice C57BL/6J mice with intratracheal (i.t.) silica (*22-27*). Hematoxylin and eosin (H & E) (Fig. 1B) and Masson trichrome staining (Fig. 1C) at 28 and 56 days post challenge revealed that i.t. silica treatment induces patchy but extensive interstitial and intra-alveolar inflammation comprised predominantly of lymphocytes and macrophages. The alveolar spaces were filled with abundant enlarged macrophages admixed with granular proteinaceous material characteristic of the pathology seen in acute silicoproteinosis. The inflammatory infiltrate was initially peribronchiolar and perivascular in distribution with progression to confluent lesions associated with collagen deposition by 56 days post treatment. Analysis of whole lung RNA isolated from silica treated mice revealed a sustained increase in expression of multiple fibrosis-related genes over the 56d time course including Tgfb1, Col1a1, Col3a1, and Fn-1 (Fig. 1D), as well as a durable increase in the hydroxyproline content of lung tissues (Fig. 1E). The pressure-volume (P-V) loop of the lungs was shifted down and to the right in silica treated mice relative to mice treated with vehicle (Fig. 1F), exhibiting a ∼30% reduction in median static compliance (0.070 ml/cmH2O vs 0.096 ml/cmH2O, respectively) (Fig. 1G). These data demonstrate that silica induces a fibrogenic program and restrictive physiologic defect that mimics silica-induced pulmonary pathologies in humans based upon gene expression, histological, biochemical, and physiological assessments.

### Single-cell analysis of the murine silicosis lung

To further examine fibrogenic programs induced by silica, we performed a longitudinal transcriptomic analysis using single-nucleus RNA-sequencing of the lungs from pre- and post-silica challenged mice. We recovered a total of 23,794 single nuclei from 12 whole lung samples across four timepoints including baseline (0 days), and 7, 28, 56 days post intratracheal (i.t.) silica). 35 unique cell states spanning epithelial, endothelial, stromal, myeloid, and lymphoid cell lineages were identified using highly and specifically expressed marker genes (Fig. 2A, 2B, and Fig. S1). Silicosis is known to produce both acute and chronic inflammation in the lung followed by progressive fibrosis. Indeed, gene set enrichment analysis revealed that inflammatory gene processes peak at Day 7 post-exposure and chronically remain higher than baseline while fibrotic gene expression processes continually increase following exposure (Fig. 2C, 2D, and E). Cell states uniquely varied both in the baseline activation of inflammatory and fibrotic gene signatures and in the subsequent temporal response following silica exposure. Many cell states were differentially abundant over time with most notable changes occurring at Day 7 post-exposure (Fig. 2C). Alveolar type I cells (AT1), bronchioalveolar stem cells, regulatory T cells, and fibrotic (Fibr-2) macrophages significantly increased in relative abundance following silica exposure while arterial, capillary, and venous endothelial cells, aerocytes, fibroblasts, mesothelial, and club cells decreased in relative abundance (adjusted p<0.05, beta-binomial test). Neuroendocrine cells demonstrated a significant yet transient increase at Day 7 post-silica exposure which may reflect a potential role in supporting acute inflammation.

**Fig. 2:**
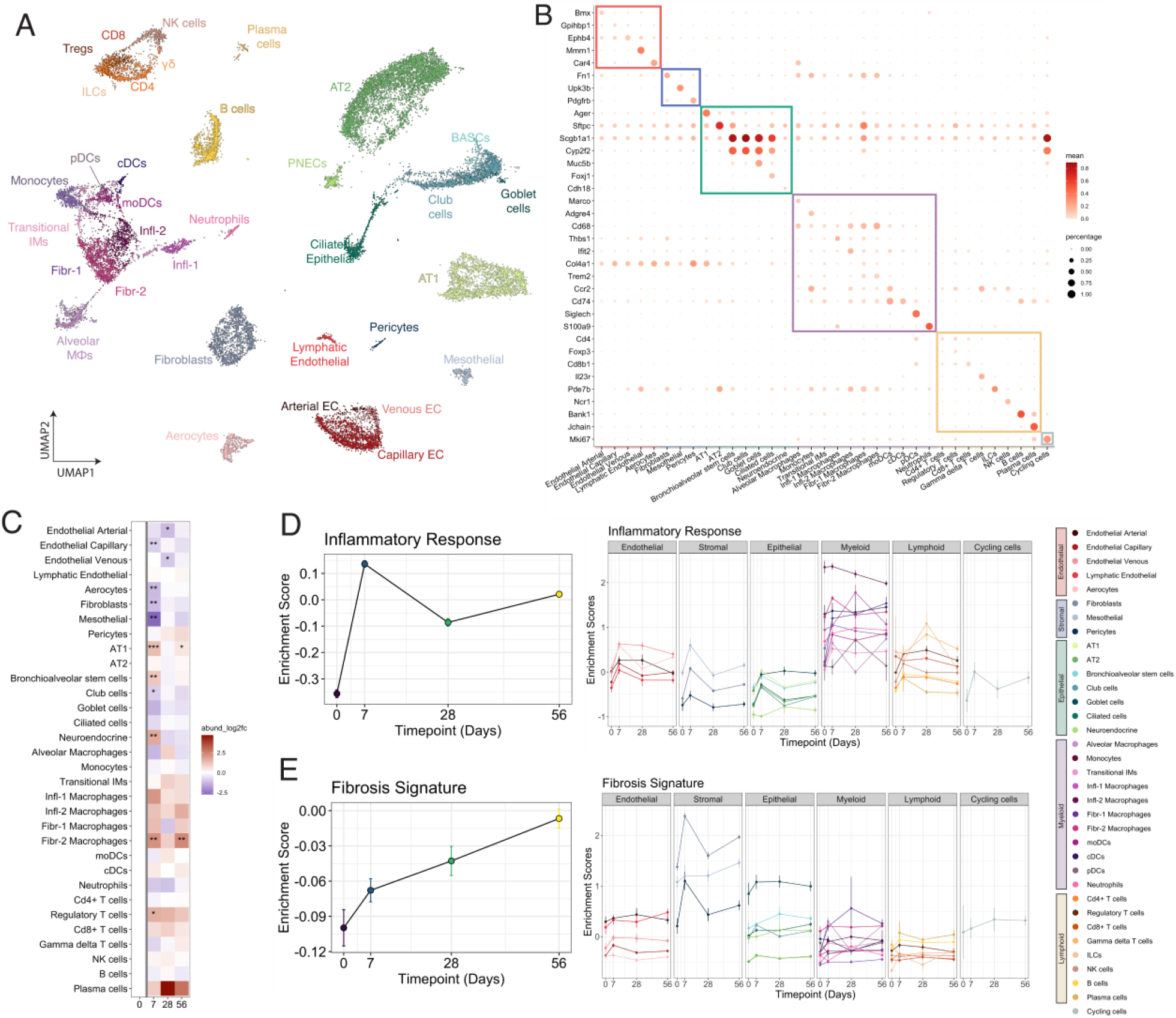
Single-nucleus RNA-sequencing depicts the global molecular dynamics in silicosis. (A) 2D UMAP representation of all annotated cell types identified. Each point represents a single cell and is colored according to fine scale annotation. (B) Selected marker genes used for broad and fine-scale cell type annotation. (C) Cell state abundance relative to Day 0 shown over time. *p<0.05, **p<0.005, ***p<0.0005; BH correction. Gene set enrichment scores for Inflammation (D) and Fibrosis (E) shown over time summarized for all cells collected from each timepoint (left panel) and colored by individual cell state (right panel).

We further interrogated the myeloid cells to identify the genetic programs activated in this lineage during silica exposure in the mouse lung (Fig. 3A). Within the myeloid cells, we identified previously described populations of alveolar (SiglecF+) and interstitial (SiglecF-) macrophages as well as populations of dendritic cells and neutrophils (Fig. 3B). Interstitial macrophages were highly abundant in the silicosis lung and demonstrated remarkable heterogeneity. Thus, interstitial macrophages (IMs) were subclustered and annotated into phenotypically descriptive groups (Fig. S2). Monocytes represented the most recently recruited cells in the lung based on high expression of C-C chemokine receptor type 2 (CCR2).

**Fig. 3:**
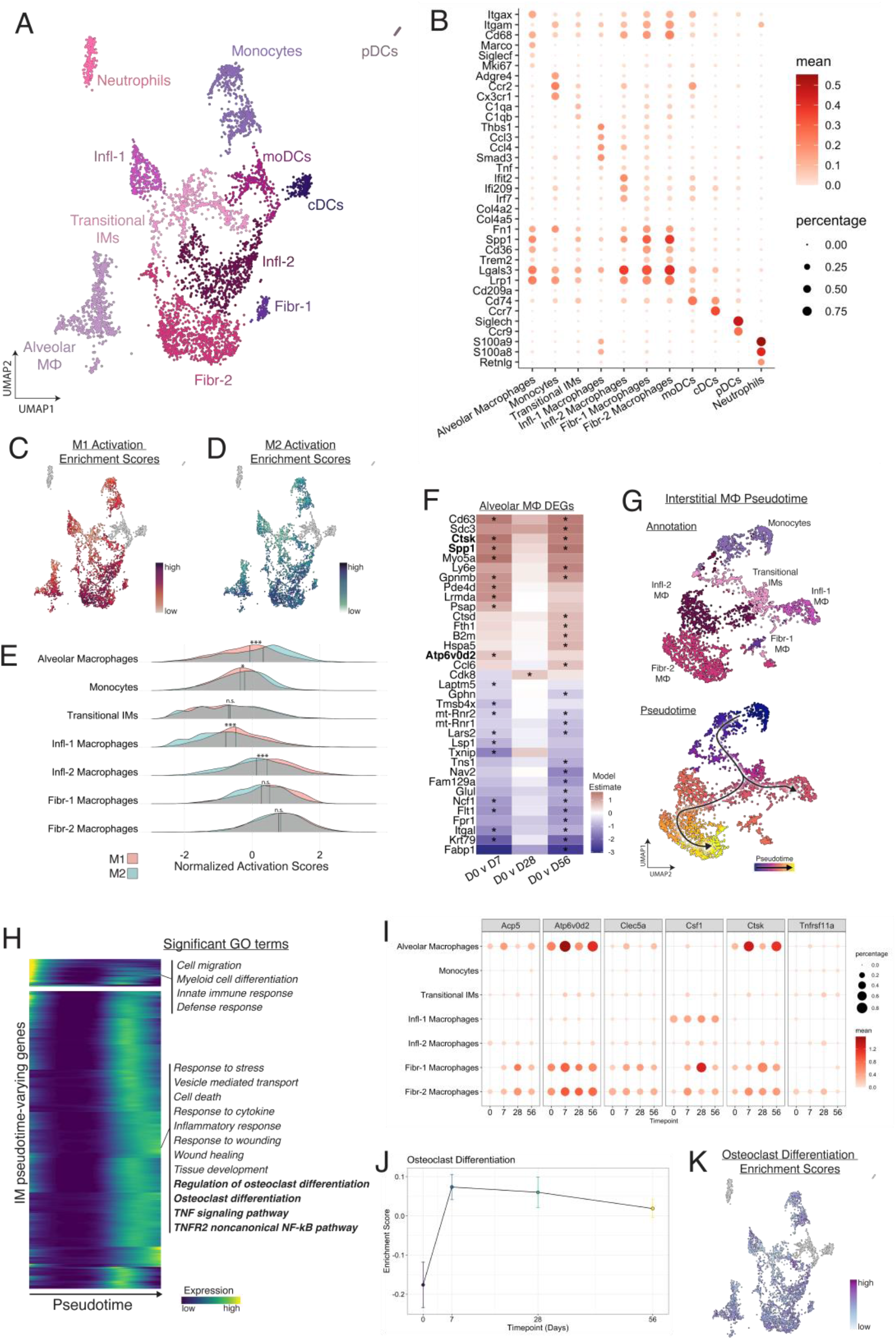
Myeloid cells show remarkable heterogeneity and activation of osteoclast-related transcriptional programs in silicosis. (A) 2D UMAP representation of myeloid cells. Each point represents a single cell and is colored according to fine scale annotation. (B) Selected marker genes used for fine-scale cell state annotation of myeloid cells. (C) M1 and (D) M2 gene sets were used to calculate an activation score for each cell. Each point is a cell colored according to the relative intensity of the signature. (E) M1 and M2 activation scores for each of the fine-scale cell state annotations was plotted in a histogram and tested for differences in mean (paired T-test). *p<0.05; BH correction. (F) Differentially expressed genes relative to Day 0 were identified for the alveolar macrophages. Genes in bold typeface are related to osteoclast differentiation and development. *p<0.05, FDR corrected. (G) 2D UMAP of interstitial macrophages colored according to fine-scale annotation (top panel) and pseudotime (bottom panel). (F) Heatmap of genes differentially expressed over pseudotime as interstitial macrophages differentiate toward pro-fibrotic phenotypes. Each row represents a gene and color represents relative intensity of expression. Select significant (q<0.05, gprofiler) representative gene ontology terms for each cluster of genes is shown on the right. (I) Expression of osteoclast genes plotted over time for each of the macrophage subsets. (J) Gene set enrichment scores for Osteoclast Differentiation were calculated for all macrophages and plotted over time. (K) Macrophages colored according to relative intensity of Osteoclast Differentiation enrichment score.

Transitional IMs express low levels of many pro-inflammatory and pro-fibrotic genes and likely represent an intermediate cell state poised to adopt a more specialized macrophage phenotype. All macrophage subsets in our analysis showed some activation of both inflammatory and fibrotic gene processes, and the traditional M1/M2 nomenclature is insufficient to describe the heterogeneity evident in our dataset (Fig. 3C-E). Thus, we annotated two populations of pro-inflammatory macrophages (Infl-1, Infl-2) and two populations of pro-fibrotic macrophages (Fibr-1, Fibr-2) based on gene expression and pathway activation differences. While both Infl-1 and Infl-2 demonstrated activation of TNF related gene expression, Infl-2 also exhibited inflammatory gene expression signaling driven by Interferon gamma (IFN-*γ*) (Fig. S2). Infl-1 peaked at Day 7 and likely represents a population of myeloid cells responsible for acute inflammation in the silicosis lung, whereas Infl-2 macrophages contribute broad inflammatory signatures evident in chronic inflammation. Infl-2 macrophages also express many pro-fibrotic genes and may serve as precursors to fibrotic macrophage cell states. Fibr-1 macrophages likely drive fibrosis due to high expression of tissue-remodeling and fibrosis-related genes including Col4a1, Fn1, and Osmr (Fig. 3B). The Fibr-2 macrophage subset likely represents end-stage pro-fibrotic macrophages in the silicosis lung due to waning expression of pro-fibrotic genes coupled with increased expression of foamy macrophage markers including Spp1, Cd36, and lipid-metabolizing genes such as Lgals3 and Lrp1 (Fig. 3B). Many pro-fibrotic genes including Spp1, Lgals3, and Lrp1 are also highly expressed in alveolar macrophages (Fig. 3B).

### Transcriptional activation of osteoclast gene programs in lung macrophages

To identify the silica-induced temporal changes in genetic programs of macrophages, we performed two analyses. First, in the SiglecF+ alveolar macrophages we identified differentially expressed genes over time. Several osteoclast marker genes were significantly increased in the alveolar macrophages of silicosis lungs compared to baseline, including Ctsk, Spp1, and Atp6v0d2 (Fig. 3F). To identify the genetic programs changing within the interstitial macrophages (SiglecF-), we performed pseudotime analysis (Fig. 3G). Pseudotime analysis linked the progression of recently recruited interstitial monocytes to a transitional state followed by pro-inflammatory and pro-fibrotic phenotypes. Genes with significantly varying expression along the pseudotime trajectory were identified, hierarchically clustered, and clusters were annotated based on gene ontology (Fig. 3H). Genes that showed increased expression later in pseudotime were significantly enriched for inflammatory response, wound healing, tissue development, osteoclast differentiation, and TNF signaling pathway (adjusted p<0.05, gProfiler). Given that osteoclastic gene expression was identified in both interstitial and alveolar macrophages, we calculated an osteoclast differentiation enrichment score using a previously published list of genes (*28, 29*) known to induce osteoclast differentiation for each macrophage subtype and plotted this over time (Fig. 3I, 3J, and 3K). The activation level of this signature increased dramatically post-silica exposure and the highest enrichment scores were present in alveolar macrophages and pro-fibrotic macrophage populations Fibr-1 and Fibr-2. Moreover, these populations expressed high levels of osteoclast markers including Acp5, Atp6v062, Clec5a, Ctsk and moderate expression of Csf1 and Tnfrsf11a (Fig. 3I).

In summary, both tissue-resident alveolar macrophages and infiltrating bone-marrow derived macrophages demonstrate robust genomic signatures of osteoclastic transformation following silica exposure. This supports the hypothesis that microenvironmental signals in the silica-challenged lung are critical in shaping pulmonary macrophage polarization states and functional phenotype.

### Osteoclast-like cells are present in the lungs of patients with silicosis and in silicosis mouse models

To validate the presence of osteoclast like cells identified by single-nucleus RNA sequencing, we performed IHC of lungs from silicotic humans and mice for two signature osteoclast proteins, TRAP and CTSK. TRAP dephosphorylates a serine on the bone ligand osteopontin (SPP1), diminishing the negative charge that facilitates binding to bone, and frees the osteoclasts to migrate to new areas of bone (*30*). Reactive oxygen species created by an iron in the catalytic site of TRAP participate in bone resorption and degradation (*31*). CTSK is a lysosomal cysteine protease participates in bone and cartilage remodeling through its capacity to degrade collagen and elastin (*32*). Despite the marked difference in disease severity, the lung tissues from both patients exhibited prominent TRAP+ and CTSK+ macrophage and MNGC accumulations within alveolar spaces surrounding small conducting airways and in focal areas of chronic interstitial inflammation adjacent to fibrotic regions (Fig. 4A and 4B). This staining pattern provides evidence of durable osteoclast-like differentiation in human silicosis. Though the life span of osteoclasts is typically measured in weeks (*33, 34*), pulmonary osteoclast-like cells are found in human silicosis patients, where the disease is typically diagnosed years after initial exposure, and persist in the silicosis lung long term in the mouse model (Fig. S3).

**Fig. 4:**
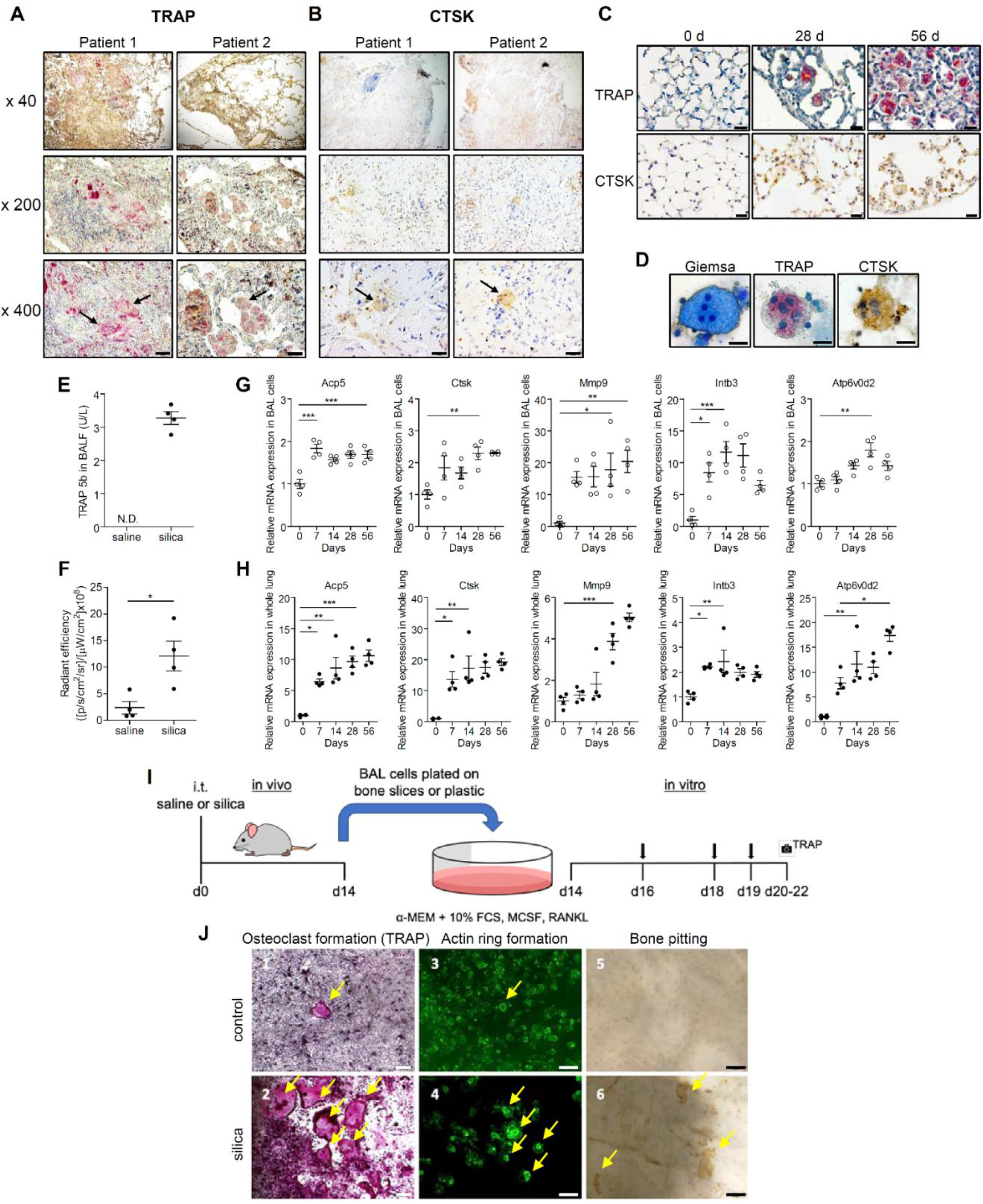
Prominent osteoclast-like phenotypes and molecular markers are evident in BAL and in whole lung of silica challenged mice. Surgical lung specimens from patients with silicosis were analyzed by TRAP staining (A), and immunohistochemical staining for CTSK (B). TRAP and CTSK positive macrophage accumulations within alveolar spaces surrounding small conducting airways and in focal areas of chronic interstitial inflammation adjacent to fibrotic regions (arrows, bottom panel) (Scale bar, 20 μm). Silica particles (5 mg, i.t.) were administered to C57BL/6J mice. (C) Paraffin-embedded lung sections collected from mice post silica treatment were stained for TRAP enzymatic activity (upper panels) and with anti-CTSK antibody (lower panels) (Scale bar, 20 μm). (D) BAL cells collected d28 post i.t. silica challenge were spun onto slides and stained with Giemsa, for TRAP enzymatic activity, and with anti-CTSK antibody. (E) The TRAP5b concentrations in BALF from saline or silica-treated mice on d28 post challenge were measured by ELISA (N = 4 mice per group) (N.D.; not detected). (F) Mice treated with saline or silica i.t. 6 days prior received the cleavage activated fluorescent cathepsin K substrate, Cat K 680 FAST by the i.t. route. The fluorescent signal was quantified 18 h later by IVIS (N = 4 mice per group). BAL cells (G, white circles) and whole lung tissue (H, black circles) were collected from mice at the indicated times in days after administration, and osteoclast-related gene expression was assessed by RT-qPCR for Acp5 (TRAP), Ctsk, Mmp9, Itgb3, and Atp6v0d2 (N = 4 mice per group). (I) Scheme for ex vivo osteoclast experiments. BAL cells isolated from naïve mice (J5), or C57BL/6J mice at day 14 post intratracheal challenge with saline (J1, J3, and J5) or silica (J2, J4, and J6) were plated on plastic (J1 and J2) or bovine bone slices (J3, J4, J5, and J6) in 96 well plates with M-CSF and RANKL. At day 6, the cells cultured on the plastic plates were stained for TRAP activity to identify multinucleated osteoclasts (J1 and J2, yellow arrows). For BAL cells plated on bone slices, actin rings were visualized by phalloidin staining (J3 and J4, yellow arrows) and resorbed bone area was visualized by peroxidase-conjugated wheat germ agglutinin/horse radish peroxidase staining after removing cells (J5 and J6, yellow arrows). Scale bars are 200 μm (J1 and J2) and 50 μm (J3, J4, J5, and J6). \**P*<0.05, \*\**P*<0.01, and \*\*\**P*<0.001

Similarly, TRAP+, CTSK+ MNGC were found in lung tissue (Fig. 4C) and bronchoalveolar lavage (BAL) (Fig 4D) of mice post silica i.t. challenge, but not at baseline. In addition, TRAP5b, the osteoclast specific isoform of the acid phosphatase ACP5 (*35, 36*), was markedly elevated in BAL fluid (BALF) of silica challenged mice but not saline control mice at d28 post instillation (Fig. 4E).

Pulmonary CTSK activity was evaluated *in vivo* by intra-tracheal administration of the CTSK cleavage activated probe, Cat K 680, to mice that had received silica vs. saline 6 days prior. Markedly increased thoracic fluorescence intensity measured by IVIS imaging 18 h post Cat K 680 administration (Fig. 4F and Fig. S4) in silica-treated mice compared to saline-treated controls is consistent with elevated pulmonary CTSK activity.

To validate the impact of silica on promoting osteoclast-like differentiation in mouse lungs, we measured RNA levels of multiple osteoclast marker genes in cells isolated by BAL and in whole lung tissue. Silica treatment increased the mRNA levels of Acp5, Ctsk, Mmp9, Intb3, Atp6v0d2, Csf1, Csf1 receptor (Csf1r), and receptor activator of nuclear factor-κB (RANK; Tnfrsf11a) (*13, 14*) in both BAL cells (Fig. 4G and Fig. S5A) and whole lung tissue (Fig. 4H and Fig. S5B). In most cases, the osteoclast genes in whole lung tissues and BAL were upregulated by day 7 and plateaued or continued to rise through day 28 or 56, providing evidence that osteoclast differentiation is an early event in silica-induced lung injury. The i.t. silica challenge also significantly increased the total cell number in BALF (Fig. S6A), due to influx of monocytes, neutrophils, and lymphocytes. MNGC were present in the BAL of silica treated mice but not saline treated controls, representing 4% of recovered cells at d7 and 6% of cells at d28 post silica treatment (Fig. 4D and Fig. S6B).

To examine whether pulmonary MNGC exhibit the signature function of osteoclasts, bone resorption, BAL cells collected from saline or silica exposed mice were plated onto plastic or bone slices and cultured in α-MEM with 10% FCS in the presence of MCSF and variable concentrations of RANKL (0-100 ng/ml) (Fig. 4I). Compared to controls (Fig. 4J1, 4J3, and 4J5), silica challenge enhanced RANKL dose-dependent formation of TRAP + multinucleated giant cells, actin rings and bone pits by isolated BAL cells (Fig. 4J2, 4J4, 4J6, and Fig. S7). BAL cells from silica exposed mice also increased the media levels of C-terminal telopeptides of type I collagen (CTX-I) when plated on bone slices, compared to BAL cells from saline exposed mice, consistent with increased bone matrix proteolytic activity (Fig. S8). Collectively these results indicate that intratracheal silica challenge of mice induces pulmonary myeloid cells to undergo osteoclastic differentiation as indicated by expression of signature osteoclast genes and proteins, formation of TRAP + MNGC, actin ring assembly and acquisition of proteolytic osteolytic functions.

### Silica-induced RANKL expression activates osteoclast-like cell formation in the lung

Osteoclastogenesis is differentially regulated by cytokines (*17, 18*). TNF-α, IL-1β, IL-6, and M-CSF stimulate osteoclast formation and function when even small amounts of RANKL are present (*17, 18*), while IL-4 suppresses osteoclast formation. Osteoprotegerin (OPG; Tnfrsf11B), acts as a soluble decoy receptor for RANKL (*13, 14, 16*), and the ratio of RANKL to OPG determines the level of free RANKL available for osteoclastogenesis (*14, 37, 38*). We found that the BALF concentrations of TNF-α, IL-1β, IL-6, and M-CSF but not IL-4 were transiently elevated by silica i.t. treatment (Fig. 5A-E). Interestingly, silica increased RANKL and OPG levels in BALF (Fig. 5F and 5G) in a staggered temporal pattern in which OPG concentrations peaked on d7, and RANKL peaked on d14 post challenge. These data indicate that the ratio of RANKL/OPG varied over time in the silica-challenged mouse lung, consistent with time-dependent differential regulation of osteoclastogenesis in the lungs of C57BL/6J mice.

**Fig. 5:**
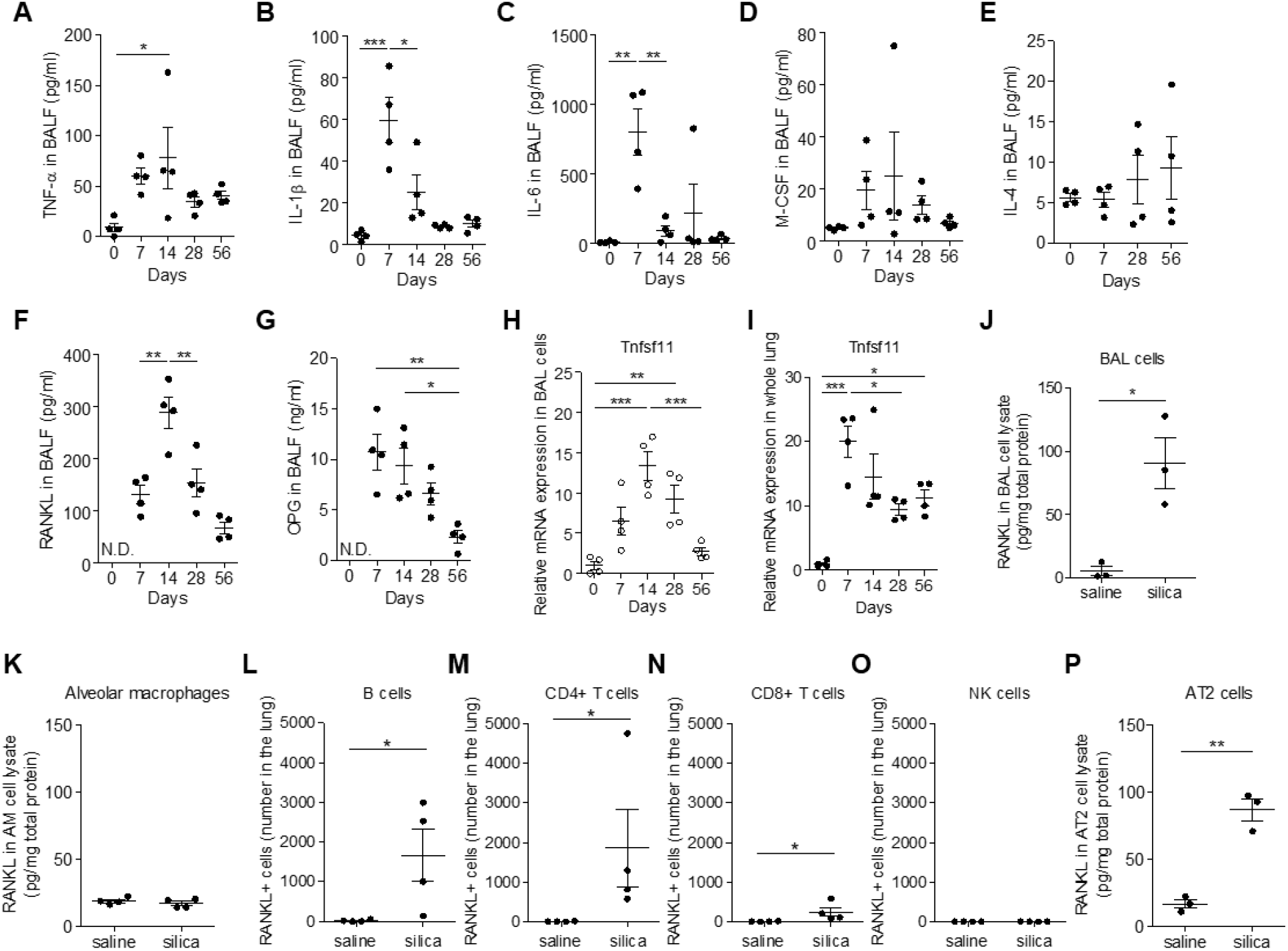
Osteoclast differentiation and activation are RANKL dependent, supported by temporal environmental cytokine expression and induced RANKL derived from lymphocytes and AT2 cells. Silica particles (5 mg) were administered i.t. into the lungs of C57BL/6J mice. Mice were sacrificed, and BALF was collected at the indicated times in days after administration. TNF-α (A), IL-1β (B), IL-6 (C), M-CSF (D), IL-4 (E), RANKL (F), and OPG (G) concentrations in BALF were measured by ELISA (N = 4 mice per group) (N.D.; not detected). (H and I) BAL cells (open circles) and whole lung homogenates (black circles) were collected at the indicated times in days after silica administration; Tnfsf11 (RANKL) gene expression was assessed by RT-qPCR (N = 4 mice per group). (J and K) RANKL in BAL cell lysates (J) and alveolar macrophage (AM) lysates (K) from silica-treated mice on d14 post silica challenge was measured by ELISA (N = 3-4 mice per group). (L-O) Whole lungs were harvested 14 days after saline or silica challenge and whole lung lymphocyte composition was determined by flow cytometry. For intracellular staining, the cells were treated with PMA, ionomycin, brefeldin A, and monensin. The number of RANKL positive and IFN-γ negative cells were determined in (L) B cells, (M) CD4+ T cells, (N) CD8+ T cells, and (O) NK cells by cell surface with intracellular staining (N = 4 mice per group). (P) RANKL in AT2 cell lysates from silica-treated mice on d14 post silica challenging was measured by ELISA (N = 3 mice per group). \**P*<0.05, \*\**P*<0.01, and \*\*\**P*<0.001.

Osteoblasts and the stromal cells that they are derived from are the classical sources of RANKL in bone (*39*), but the origin of RANKL in the lung is unknown. We found that silica treatment upregulated RANKL gene expression in both BAL cells and whole lung homogenates (Fig. 5H and 5I). Silica treatment increased RANKL protein expression in BAL cell lysate, but not in the purified AM cell lysate (Fig. 5J and 5K). Activated lymphocytes have been reported to be a source of RANKL in inflammatory responses (*40, 41*) and silica exposure increased the number of lymphocytes in BALF (Fig. S6). Therefore, RANKL expression in pulmonary lymphocytes was assessed by flow cytometry. Silica exposure induced low level but detectable cell surface and intracellular RANKL expression in pulmonary B cells, CD4+ T cells, and CD8+ T cells (Fig. 5L-N). In contrast, RANKL was not induced in natural killer (NK) cells (Fig. 5O). Furthermore, RANKL expression in alveolar type II (AT2) cells isolated from silica exposed mice was evaluated given that AT2 cells are known to secrete various cytokines (*42, 43*).

RANKL protein expression was increased in cell lysates of AT2 cells isolated from mice d14 silica mice compared to d14 saline challenged mice (Fig. 5P), and RANKL mRNA expression was increased in isolated, cultured rat AT2 exposed to silica ex-vivo (Fig. S9). Collectively, these data indicate that intratracheal silica treatment induces RANKL expression in pulmonary lymphocytes and AT2 cells, which likely represent the sources of the cytokine that induce pulmonary osteoclastogenesis of recruited monocytes and resident macrophages.

### Silica-induced osteoclast-like differentiation is RANKL-dependent

To examine the role of RANKL in osteoclast-like differentiation in the silica-challenged lung, we investigated whether a neutralizing anti-mouse RANKL monoclonal antibody (mAb, clone; IK22-5) (*15, 44-47*) delivered by the intraperitoneal (i.p.) route three times weekly attenuates silica-induced osteoclastogenic programs (Fig. 6A). Anti-RANKL mAb treatment suppressed silica-induced expression of osteoclast-related genes Acp5, Ctsk, Mmp9, and Atp6v0d2 in both BAL cells (Fig. 6B) and in whole lung homogenates (Fig. 6C) compared to control (Ctrl) IgG. Anti-RANKL also reduced TRAP enzymatic activity in tissues sections compared to Ctrl IgG, especially within MNGC (Fig. 6D). Expression of CTSK determined by IHC was also attenuated by anti-RANKL treatment both in whole lung tissues and in MNGC (Fig. 6D). Anti-RANKL mAb treatment decreased the levels of TRAP 5b in BALF compared with Ctrl IgG (Fig. 6E). Collectively, these data indicate that silica treatment activates osteoclast-like differentiation in the lung in a RANKL dependent manner.

**Fig. 6:**
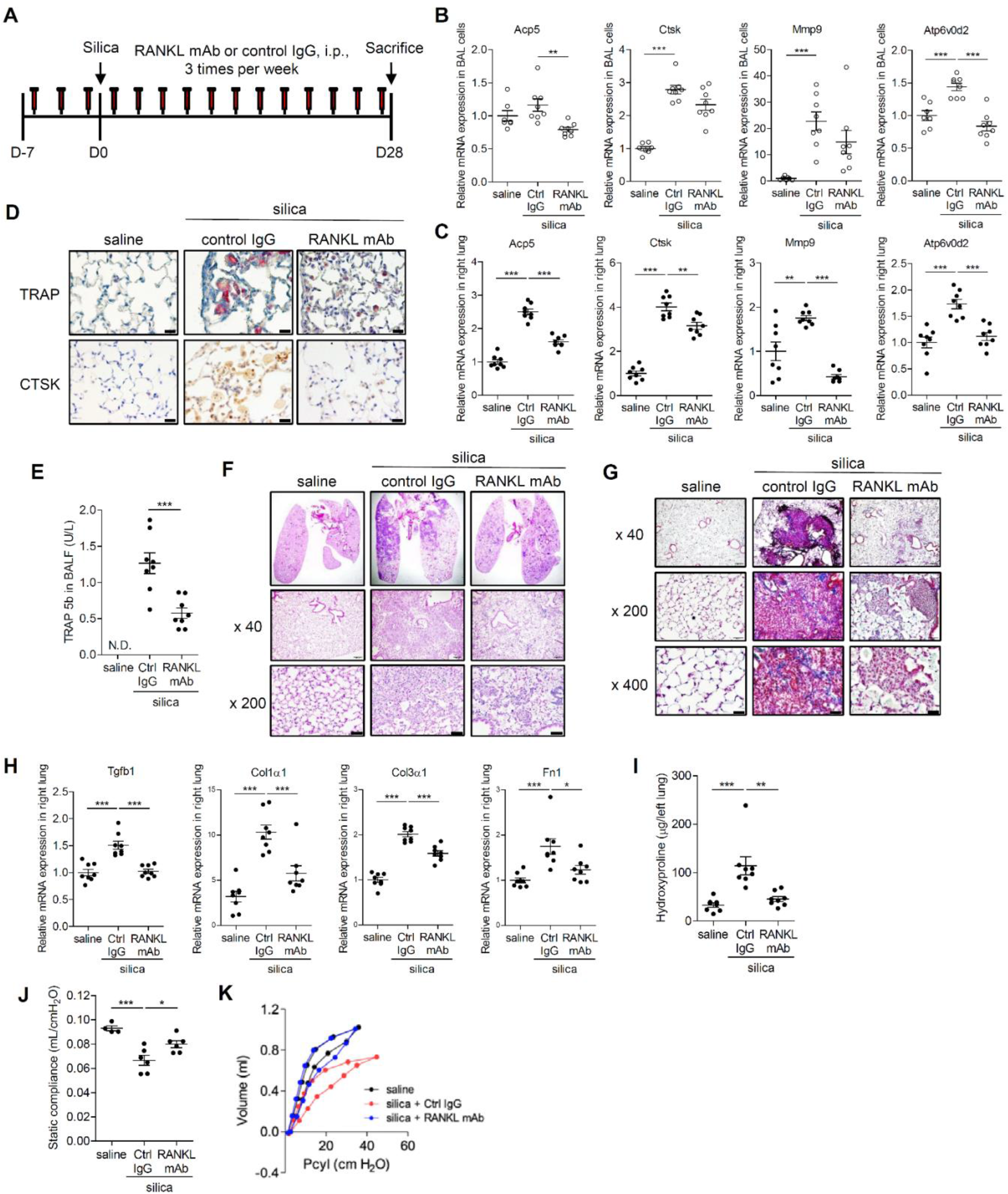
Inhibiting the RANKL dependent differentiation of pulmonary osteoclast like cells ameliorates pulmonary fibrosis. (A) Schematic depicting study design for anti-RANKL mAb studies. C57BL/6J mice treated with anti-mouse RANKL mAb or Rat IgG2a isotype control (Ctrl IgG) (0.25 mg/mouse, i.p., 3 times per week) were challenged with silica particles (5 mg) and sacrificed at 28 days later. (B and C) BAL cells (B, open circles) and whole right lung tissues (C, black circles) were collected and osteoclast-related gene expression was assessed by RT-qPCR for Acp5 (TRAP), Ctsk, Mmp9, Atp6v0d2 (N = 7-9 mice per group). (D) Paraffin-embedded lung sections were stained for TRAP enzymatic activity (upper panels) or with anti-CTSK (lower panels) (N = 6 mice per group) (Scale bar, 20 μm). (E) The levels of TRAP 5b in BALF were measured by ELISA (N = 7-9 mice per group) (N.D.; not detected). Paraffin-embedded lung sections were stained with (F) H & E and (G) Masson’s trichrome reagent (N = 6 mice per group) (Scale bar, 20 μm). (H) Right lungs were harvested and fibrosis-related gene expression was assessed by RT-qPCR for Tgfb1, Col1α1, Col3α1, and Fn1 (N = 7-9 mice per group). (I) Hydroxyproline levels in left lungs were quantified (N = 7-9 mice per group). Pulmonary function tests were performed. (J and K) The representative curves of pressure in the cylinder (Pcyl)/volume from the three groups and compliance measurements were made using the forced oscillation method and plotted as described in Materials and Methods (N = 4-6 mice per group). \**P<*0.05, \*\**P<*0.01 and \*\*\**P<*0.001.

### Anti-RANKL mAb suppresses silica-induced pulmonary fibrosis

We postulated that tissue injury from osteoclast-derived matrix degrading proteases and hydrochloric acid are key drivers of progressive pulmonary fibrosis in silicosis. Multiple histologic, immunohistochemical, and physiologic approaches were applied to determine if anti-RANKL mAb attenuates silica-induced pulmonary disease. H & E (Fig. 6F) and Masson trichrome staining (Fig. 6G) revealed that silica dependent pulmonary inflammation and fibrosis was significantly attenuated by 5 weeks of anti-RANKL antibody treatment as compared to Ctrl IgG treatment. Silica-induced upregulation of pulmonary expression of fibrosis-related genes including Tgfb1, Col1α1, Col3α1, Fn1 (Fig. 6H) and the accumulation of hydroxyproline in lung tissues were partially blocked by anti-RANKL mAb treatment (Fig. 6I). In addition, the P-V curve of silica-treated mice receiving Ctrl IgG was shifted down and to the right, consistent with reduced compliance, whereas the position of the P-V curve of silica-treated animals receiving anti-RANKL mAb partially corrected toward baseline, consistent with partial rescue of silica-related reduction in compliance (Fig. 6J and 6K). These data implicate pulmonary osteoclasts like cells in the pathogenesis of silica induced pulmonary fibrosis.

## DISCUSSION

Silicosis is an intractable and inexorably progressive form of pulmonary fibrosis that remains a global occupational threat. Myeloid cells, including monocytes and alveolar macrophages, play a central role in disease pathogenesis including the recruitment of lymphocytes and other inflammatory cells and signaling to stromal cells that express excess extracellular matrix and distort the lung architecture (*11, 48*), but the mechanisms involved are incompletely understood. Scavenger receptors with collagenous domains expressed on the macrophage surface recognize silica (*12*) and mediate destructive remodeling in the lung (*4, 12, 49, 50*) through elaboration of reactive oxygen species and matrix degrading enzymes (*50*).

Phagocytosis of silica activates the inflammasome and induces IL-1β and other cytokines which play important roles in subsequent inflammation and fibrosis (*4, 49, 51, 52*), Recent studies demonstrate that silica-induced foam cell formation in alveolar macrophages leads to release of TGF-β1 and promotes pulmonary fibrosis (*53*). The pathophysiologic relevance and relative contribution of each of these fibrogenic processes in the development of the silicotic lesion in vivo remains unclear, however, as do the final effectors of tissue damage.

We employed single nucleus RNA-sequencing on longitudinally collected whole lung samples to explore novel mechanisms of silica induced fibrosis. We identified a multitude of distinct cell states that temporally contribute to the inflammatory and fibrotic pathways evident in the silicotic lung. We detected activation of an osteoclast program in both tissue resident and recruited monocytes, indicating that the shift was lineage-independent and driven by tissue specific microenvironmental factors. We validated the presence of osteoclast like cells and osteoclast differentiation in human and mouse lung tissues using rtPCR, immunohistochemistry, biomarker analysis, and osteoclast functional assays (multinucleation, bone pitting, actin ring formation, TRAP and CTSK activity). We determined that silica challenge induced regional expression of the signature osteoclastogenic cytokine, RANKL, and identified the lung cell compartments that it was derived from, lymphocytes (*40, 41*) and AT2 cells. Collectively, these data suggest aberrant induction of a gene program in silica exposed lung macrophages drives a pathogenic response to lung injury.

Because silica particles are non-dissolvable and remain in the lung indefinitely, they may stimulate RANKL expression in a persistent manner and contribute to ongoing differentiation of pulmonary osteoclast-like cells. The long-term persistence of osteoclast-like cells after silica challenge is more consistent with the durable temporal pattern of RANKL expression than the transient expression of the cytokines that have been implicated in inflammasome associated inflammatory osteolysis, TNF-α, IL-1β, and IL-6. Our longitudinal data suggest that RANKL is the major osteoclastogenic stimulus in the lung following silica challenge. Indeed, disruption of the RANKL/RANK axis with a monoclonal antibody attenuated osteoclastogenesis and the histological, immunohistochemical, genetic, biochemical, and physiological hallmarks pulmonary fibrosis, suggesting RANKL as a promising candidate for future disease intervention.

It remains possible that anti-RANKL treatment dampens silica-induced pulmonary fibrosis through downregulation of an inflammatory pathway that is independent of osteoclast formation. The RANKL/RANK axis plays important roles in the immune cell maturation and differentiation during development, including in lymph-node formation, lymphocyte differentiation, dendritic cell survival, T-cell activation, and tolerance induction (*18*) but is generally believed to play a relatively minor role in inflammation in the adult organism. Indeed, the anti-RANKL monoclonal antibody (*54, 55*), denosumab, that is widely used as an osteoporosis therapy is not known to have significant immunosuppressive or anti-inflammatory properties.

Osteoclasts have not been previously reported to play role in lung homeostasis or pathophysiology. The first evidence for a bonafide pulmonary role for osteoclasts emerged from our study of pulmonary alveolar microlithiasis (PAM), a rare, autosomal recessive and often fatal disorder of pulmonary phosphate metabolism caused by mutations in the epithelial sodium phosphate co-transporter, NPT2b, that lead to accumulation of calcium phosphate crystals in the alveolar lumen (*56, 57*). We found that epithelial deletion of NPT2b in mice results in a progressive pulmonary process characterized by diffuse alveolar microlith accumulation (*58*), development of pulmonary fibrosis, and, surprisingly, accumulation of surfactant proteins and phospholipids consistent with alveolar proteinosis. In both PAM patients and our NPT2b^-/-^ mouse model we discovered that osteoclastic transformation of alveolar macrophages (AM) and alveolar monocytes are key components of the host defense against microliths in the PAM lung that mediate clearance of the particle (*59*). In light of our findings with silica, we postulate that polarization of alveolar macrophages or recruited monocytes toward an osteoclast phenotype and away from a surfactant catabolizing phenotype is a stereotypical response to exposure to particles, including microliths, silica and perhaps others, and that the acid and enzyme products of pulmonary osteoclasts contribute to pulmonary fibrosis.

Osteoclastogenesis has rarely been reported in compartments other than bone, and it remains unclear whether the pulmonary osteoclast-like cells in the silicosis model represent the fully differentiated osteoclasts found in bone. Yet these ectopic cells demonstrate the most specific distinguishing features of true osteoclasts including formation of actin rings, and the capacity to degrade the mineral and matrix components of bone. Whether better described as osteoclasts or osteoclast-like, these cells are clearly differentiating along an osteoclast pathway, and represent an interesting new pulmonary myeloid cell phenotype of potential pathophysiologic significance.

In summary, we find that pulmonary silica exposure incites progressive fibrosis in part via RANKL-dependent osteoclastic transformation of pulmonary myeloid cells, likely through their sustained elaboration of matrix degrading proteases and hydrochloric acid. These findings showcase the utility of whole tissue single nucleus sequencing combined with multiple modes of biological validation to identify both a novel mechanism of pulmonary fibrosis and a promising new therapeutic target for silicosis, the pulmonary osteoclast-like cell. Though osteoclastic transformation of macrophages is a known component of bone homeostasis in adult organisms, our study demonstrates that activation of a routine genetic program in an atypical location can produce a pathological result. This study illustrates how collecting molecular measurements of all genes in all cell types at once in an authentic disease model is a powerful approach to understanding the pathobiology of the disease and identifying new molecular targets.

## MATERIALS AND METHODS

### Study Design

Silicosis is a life-threatening, progressive lung disease limited by a poor understanding of pathophysiology and a lack of effective treatments. We explored the mechanisms of silica-induced pulmonary fibrosis in a mouse model using multiple modalities including whole-lung single-nucleus RNA sequencing. Silica particles were administered via the i.t. route into the lungs of C57BL/6 mice. Lung fibrosis in tissues from silicosis patients and silica-challenged mice was characterized, using fibrosis-related gene expression quantified by RT-qPCR, hydroxyproline measured in lung homogenates, and lung compliance measured using oscillatory impedance. To explore novel mechanisms of silica-induced fibrosis, single nucleus RNA-sequencing was conducted on longitudinally collected whole lung samples. To validate these findings, the presence of osteoclast-like cells and osteoclast differentiation in human and mouse lung tissues was interrogated with RT-qPCR, immunohistochemistry, biomarker analysis, and osteoclast functional assays. Cytokines known to participate in osteoclast differentiation were measured in BALF by ELISA. The finding that silica exposure induced regional expression of the signature osteoclastogenic cytokine, RANKL, led to a search for the source. Increased RANKL protein expression was identified after silica challenge in isolated AT2 and lymphocytes by ELISA and flow cytometry. The propensity of cultured BAL cells isolated from mice before and after silica challenge was tested by determining the threshold of RANKL concentration required for robust, multinucleated osteoclast formation. Fibrosis assessed silica-challenged mice treated with anti-RANKL by the i.p. route using histology, RT-qPCR, hydroxyproline assay, and pulmonary function testing. Because the silica-challenged animals were easily distinguished from controls based on the presence of particles in lung tissue, randomization and blinding were not used for experiments with animals, but mice were age- and -sex matched for all studies. The experimental procedures were approved by the Institutional Animal Care and Use Committee at the University of Cincinnati.

### Animals

C57BL/6J mice were obtained from Jackson Laboratory (Bar Harbor, ME). Sprague-Dawley rats used for alveolar type II cell isolation were obtained from Charles River Laboratory (Wilmington, MA). For terminal experiments, mice or rats were euthanized by i.p. injection of Euthasol (Henry Schein, Melville, NY). Chow that contained 0.7% phosphate, 16.3% protein, 66.3% carbohydrate, 5.0% fat, 1.2% calcium and vitamin D3 (2900 IU/Kg) was prepared by Harlan Laboratories (Madison, WI) and purchased from Harlan Sprague Dawley (Indianapolis, IN, USA). All animals were maintained in a specific pathogen–free facility and were handled according to a University of Cincinnati Institutional Animal Care and Use Committee approved protocol and National Institutes of Health guidelines.

### Histology

Archived, anonymized surgical lung specimens from human patients with silicosis were kindly provided by Ciara Shaver, M.D. Vanderbilt University Medical Center (Nashville, TN). Lung tissues were fixed with 10% buffered formalin phosphate, embedded in paraffin, and stained with H & E. Masson’s trichrome staining was used to identify collagen deposition (Newcomer Supply, Middleton, WI).

### In vivo exposure to silica

Silica particles (Sigma Aldrich, St. Louis, MO, particle size: 80% between 1 and 5 µm) were boiled in 1N HCl for 1 h, washed with dH2O, and dried at 100°C. The particles were then heat sterilized at 200°C for 2 h and suspended in sterile saline. The endotoxin content in the silica particles was <1.0 pg/μg of silica as determined using the LAL Chromogenic Endotoxin Quantitation Kit (Thermo Scientific, Rockford, IL) according to the manufacturer’s instructions. For i.t. silica administration, C57BL/6J mice were anesthetized with isoflurane and suspended by their incisors in the supine position on a procedure board at a 45° angle. The glottis was visualized by retraction of the tongue and illuminated with a fiberoptic thread. A 22-gauge angiocatheter was advanced into the trachea under direct visualization, and after confirming correct placement by expansion of the thorax upon delivery of air though the catheter, 2.5 or 5 mg of silica in 100 μl of saline was injected into the lung. For oropharyngeal (o.a.) silica administration, each mouse was anesthetized with isoflurane and suspended by a steel wire on a procedure board at a 60°C angle by the incisor teeth. The mouth was opened, the tongue was pulled forward and 5 mg of silica in 50 μl of saline was placed at the base of the tongue. Once the slurry was aspirated into the lungs with inspiration, the tongue was released.

### Immunohistochemistry and immunocytochemistry

Immunohistochemical analysis was performed on formalin-fixed, paraffin embedded lung specimens. Tissue sections were deparaffinized, rehydrated with dH2O, subjected to antigen retrieval with citrate buffer, and blocked with normal serum. For TRAP staining, tissue sections were incubated with pre-warmed (37°C) TRAP staining mix (50 mM MES, pH 4.8, containing 50 mM L-(+) tartaric acid, 0.3 mM naphthol AS-MX phosphate, and 1.5 mM Fast Red Violet LB Salt (Sigma Aldrich)) for 4 h and counterstained with hematoxylin for 5 sec. For immunocytochemistry, cells were cytospun onto glass slides (700 rpm, 5 min), dried overnight, fixed with 3.7% neutral buffered formalin. Specimens were incubated with anti-cathepsin K antibody (polyclonal, 1:750) obtained from Abcam (Cambridge, UK), followed by horseradish peroxidase (HRP) linked anti-rabbit secondary antibody (Cell Signaling Technology, Beverly, MA). The DAB substrate kit (Thermo Scientific) was used for color development and samples were counterstained with hematoxylin for 5 sec.

### Sci RNA-Seq Library Generation

Single nuclei RNA-seq was felt to be preferable to single cell RNA-seq to avoid exclusion of multinucleate osteoclast like cells. Single nuclei RNA-Seq data was prepared using the 2-level workflow for sci RNA-Seq (*60*). The protocol was modified to use the following RT incubation temperatures, instead of a 55°C /5 minute incubation: 2 minutes at each of 4°C, 10°C, 20°C, 30°C, 40°C, 50°C followed by 10 minutes at 53°C and 15 minutes at 55°C. Samples were prepared using 6 plates of RT indices (for 576 indices used) divided evenly between the 12 samples. 150 nuclei in total were sorted into each well of the 4 PCR plates that were prepared. Libraries were sequenced using an Illumina Nextseq 550.

### Sequencing data processing and analysis

Raw sequencing output was processed as previously described(*61*) and analysis was performed using Monocle 3 (v1.2.9) (*62*). Cells were filtered for quality control based on the following thresholds: >100 UMIs, <10% mitochondrial RNA, <0.2 Scrublet (*63*) doublet score. For visualization, we performed dimensionality reduction using the first 100 principal components, followed by 2D UMAP projection. We hierarchically annotated the data (into broad and fine cell-states) using Louvain clustering with varying levels of resolution in combination with marker genes identified from literature.

### Differential Abundance testing

Cell numbers were collapsed per sample according to the fine cell state annotation. Cell numbers were size factor normalized to correct for different cell numbers recovered from samples. A beta-binomial test with Benjamini-Hochberg correction for multiple hypotheses was used to test for differences in cell state abundance for each timepoint compared to baseline (Day 0). A corrected p-value less than 0.05 was considered significant.

### Gene signature analysis

Gene sets were acquired from MSigDB (*29*) (Hallmarks Inflammation Gene Set, GO_Osteoclast_differentiation), Aran et al 2017 (*64*) (M1/M2 signatures), and Wang et al(*65*) (Fibrosis signature). For each gene, we translated the human gene set to mouse orthologs using the gorth function from the gprofiler2 package in R. For M1 and M2 analyses, only genes unique to one of the lists was used. A summary score of gene set activation was calculated per cell using the aggregate_gene_expression function in Monocle3 (*62*). Scores were calculated using log-transformed expression values and were normalized to a scale of -3 to 3.

### Differential Expression analysis

To test for differences in gene expression over time in the sequencing experiment, we used a linear mixed effect model with splines. Significant differentially expressed genes were annotated with gene ontology (GO) function terms using g:Profiler. Terms with p<0.05 corrected for multiple hypothesis testing with the default g:SCS method were considered significant.

### Pseudotime analysis

We calculated pseudotime for the interstitial macrophages to capture pathway activation changes associated with macrophage polarization and plasticity. Pseudotime was calculated using learn_graph in Monocle3 (*62*) with default parameters except for use_partitions = FALSE. The root node was designated based on end with highest number of cells collected from Day 0. Genes, expressed in at least 100 cells, that vary along the pseudotime trajectory were identified using linear model with natural splines (df = 3). Differentially expressed genes were selected based on q_value<0.05. Significant differentially expressed genes were annotated with gene ontology (GO) function terms using g:Profiler. Terms with p<0.05 corrected for multiple hypothesis testing with the default g:SCS method were considered significant.

### Fluorescent live imaging

Cat K 680 FAST Fluorescent Imaging Agent (PerkinElmer Inc., Hopkinton, MA) was used to *in vivo* detect cathepsin K activity in the lung. C57BL/6J mice that had been intra-tracheally challenged with silica or saline one week prior by the i.t. route were shaved around the chest prior to imaging, and 1 nmol of Cat K 680 FAST probe was administered by the i.t. route. The fluorescence signal emanating from the chest was monitored using fluorescent live imaging system (IVIS Spectrum, PerkinElmer Inc.) at 18 h post-injection and fluorescence intensity was analyzed as previously described (*66, 67*).

### Lung physiological measurements using SCIREQ Flexivent System

C57BL/6J mice were anesthetized with isoflurane or ketamine and intubated through a tracheostomy with a metallic angiocatheter. Lung compliance and pressure volume characteristics were measured using oscillatory impedance (Flexivent, version 5.1, SCIREQ Scientific Respiratory Equipment Inc. Montreal, Canada) and plotted using GraphPad Prism (Ver.5.03., GraphPad Software, San Diego, CA) as previously reported (*58, 68*).

### Osteoclast formation, bone resorption pit assay, actin ring assembly

BAL cells were collected d14 post i.t. saline or silica challenge and plated on plastic or bovine bone slices (Immunodiagnostic Systems, UK) in 96-well plates (2.0 × 10^4^ cells/well) in α-minimum essential medium (α-MEM; Gibco) containing 10% FBS in the presence of various concentrations (0-100 ng/ml) of recombinant murine RANKL (PeproTech) and 1:40 dilution of CMG12-14 (murine M-CSF producing cell line (*69*)) conditioned medium (equivalent to 30 ng/ml of recombinant human M-CSF). Medium was changed on day 2, 4, and 5. Cells plated on plastic were fixed in formalin and stained for TRAP activity after 6 days in culture. For actin ring staining, cells on bone slices were fixed on day 6 in formalin and permeabilized in 1% Triton X-100, rinsed with PBS, and stained with AlexaFluor 488-phalloidin (Invitogen) (*70*). For the bone pitting assay, cells on bone slices were removed by scrubbing the bone surface with toothbrush, and resorption pits were visualized by incubation with 20 µg/ml peroxidase-conjugated wheat germ agglutinin and staining with 3,3’-diaminobenzidine (Sigma-Aldrich) (*69, 70*). The levels of CTX-I in culture medium over bone slices were measured using the CrossLaps^®^ for culture (CTX-I) ELISA (Immunodiagnostic Systems) (*71*).

### Anti-RANKL monoclonal antibody treatment

C57BL/6J mice were treated with an anti-mouse RANKL mAb (clone IK22-5, Bio-X-Cell, West Lebanon, NH) or a Rat IgG2a isotype Control IgG (clone 2A3, Bio-X-Cell) (0.25 mg/mouse, i.p., three times per week) one week prior to i.t. or o.a. challenge with 5 mg of silica particles. After the silica administration, mice continued to be treated with the anti-RANKL mAb or a Rat IgG2a isotype IgG three times per week, and then sacrificed 28 days post silica administration, followed by assessments by RT-qPCR, immunochemistry, histology, hydroxyproline assessment of lung tissues and BAL cell. Some mice underwent pulmonary function testing as described above.

### Data access

We used the genes annotated to “osteoclast differentiation” from gene ontology (downloaded from MSigDB website link: https://www.gsea-msigdb.org/gsea/msigdb/human/geneset/GOBP_OSTEOCLAST_DIFFERENTIATION.html). Sequencing data will be submitted to GEO following acceptance of the manuscript.

### Statistical analysis

Statistical analyses were performed using GraphPad Prism (version 5.03, GraphPad Software). Differences between two groups were compared using the Student’s *t* test, an unpaired *t* test with the Welch correction, or Mann-Whitney U test. In experiments in which more than two groups were involved, one-way analysis of variance (ANOVA) was used followed by Bonferroni’s multiple comparisons test. Differences were considered significant at p < 0.05.

## Supplementary Materials

### Materials and Methods

Fig. S1. Fine annotation of single-nucleus RNA-sequencing data.

Fig. S2. Analysis of gene expression and pathway activation differences to characterize myeloid cell states.

Fig. S3. Silica-induced osteoclast-like cells persist up to 1 year post silica i.t. challenge

Fig. S4. Intratracheal silica challenge induces CTSK enzyme activity in the lungs of mice.

Fig. S5. Osteoclast-related gene expression in BAL cells and lung tissue from silica exposed mice.

Fig. S6. BAL cell numbers and differential in i.t. silica challenged mice.

Fig. S7. Intratracheal silica challenge enhances osteoclast formation, bone pitting and actin ring assembly is cultured BAL cells.

Fig. S8. Intratracheal silica challenge enhances bone matrix degradation by BAL cells cultured on bone slices.

Fig. S9. Silica particles induce RANKL expression in primary rat AT2 cells.

Table S1. Primers for quantitative RT-PCR

## Author contributions

Y.H. designed and performed experiments, analyzed data, and wrote the manuscript. J.F conducted and analyzed single nucleus RNA sequencing experiments and wrote the manuscript. Y.T. developed the silicosis model mouse and performed bone pitting assay and ELISA. Y.U. developed the silicosis model mouse and performed IVIS assays. DFR, CW, SS performed single nucleus RNA sequencing experiments. L.B.P. performed histological and pulmonary physiological experiments. N.M.N. designed experiments and analyzed data. C. M. S. shared surgical lung specimens from human silicosis patients. H.W. conducted experiments with isolated rat and mouse AT2 cells. J.C.G. assisted with single nuclei analysis and flow experiments. A.R.O. analyzed flow cytometry data. J.J.Y provided oversight for the IVIS assay. E.J.K. wrote the manuscript. S.L.T. analyzed data and provided osteoclast expertise. K.W.B. performed pathological assessments and wrote the manuscript. CT oversaw single nucleus RNA sequencing experiments and wrote the manuscript. FXM developed the concept, designed experiments, analyzed data, and wrote the manuscript. All co-authors read and edited the manuscript.

## Data and materials availability

All data are available in the main text or the supplementary materials.

