## Supplemental Materials for "Pulmonary osteoclast-like cells in silica induced pulmonary fibrosis"

**This file includes:**

Materials and Methods

Fig. S1. Fine annotation of single-nucleus RNA-sequencing data.

Fig. S2. Analysis of gene expression and pathway activation differences to characterize myeloid cell states.

Fig. S3. Silica-induced osteoclast-like cells persist up to 1 year post silica i.t. challenge

Fig. S4. Intratracheal silica challenge induces CTSK enzyme activity in the lungs of mice.

Fig. S5. Osteoclast-related gene expression in BAL cells and lung tissue from silica exposed mice.

Fig. S6. BAL cell numbers and differential in i.t. silica challenged mice.

Fig. S7. Intratracheal silica challenge enhances osteoclast formation, bone pitting and actin ring assembly is cultured BAL cells.

Fig. S8. Intratracheal silica challenge enhances bone matrix degradation by BAL cells cultured on bone slices.

Fig. S9. Silica particles induce RANKL expression in primary rat AT2 cells.

Table S1. Primers for quantitative RT-qPCR

References (72, 73)

### MATERIALS AND METHODS

**Cytology.** For cytology, BAL cells were spun onto glass slides (700 rpm, 5 min), dried overnight, and incubated with Giemsa solution (Sigma Aldrich) for 1 min and washed with $dH_2O$ for 5 min, three times.

**BAL cells and fluid collection.** Mice were sacrificed and tracheostomy was performed. A plastic cannula was inserted through the incision and the lung was flushed with 1 ml of saline and repeated five times. The total volume recovered was approximately 4 ml. The BALF was centrifuged at 500 × *g* for 10 min to pellet the cells. Gene expression in BAL cells was assessed by quantitative reverse transcription PCR (RT-qPCR), and the cytokines in the supernatant from the first lavage cycle were quantified by enzyme-linked immunosorbent assay (ELISA).

**Preparation of RNA and RT-qPCR.** Total RNA was isolated from murine BAL pellet and murine whole lung tissues using RNAzol RT (Molecular Research Center, Cincinnati, OH), according to the manufacturer's instructions. cDNA was synthesized using the High-Capacity cDNA Reverse Transcription Kit (Applied Biosystems, Waltham, MA). RT-qPCR was performed using a SYBR Green Master Mix (Applied Biosystems) with primer pairs for fibrosis-related and osteoclast-related genes, as well as β-actin (Actb) and actin α-2 smooth muscle (Acta2) as internal controls. The nucleotide sequences of the primer pairs are included in the Table S1.

**Measurement of hydroxyproline content in the mouse lung.** For hydroxyproline measurements, lung tissues were homogenized with 1 ml of $dH_2O$ per 100 mg of tissue. Equal volumes of 12 M HCl and lung homogenate were mixed, and the samples were hydrolyzed at 120°C for 3 h and centrifuged at 10,000 x *g* for 3 min. The supernatant was collected and hydroxyproline in the supernatant was determined using

Hydroxyproline Assay Kit (Sigma Aldrich), according to the manufacturer's instructions.

**Lung Dissociation and Nuclei Fixation.** Snap-frozen mouse lung tissue (93-223 mg) was dissociated using a Gentle MACS tissue dissociator using "C" dissociation tubes in 5 ml of ice-cold lysis/fixation buffer (10 mM NaCl, 10 mM sodium phosphate pH 7.2, 3 mM $MgCl_2$, 5% glutaraldehyde, 10 mM vanadyl ribonucleoside complex, 0.1% Triton X-100, 1% diethyl pyrocarbonate, 0.00015% polyvinyl sulfonic acid (Sigma Cat. 278424)). Tissue was dissociated using the "Mouse Spleen 1" program (gentleMACS Dissociator, Miltenyi Biotec) for 60 seconds, then filtered using a 70 µM cell strainer. The strainer was washed with an additional 5 ml lysis/fixation buffer, then nuclei were fixed at 4°C for 15 minutes and pelleted by centrifugation at 500 RCF, 4°C, for 8 minutes. Supernatant was discarded and nuclei were resuspended in 1 ml of nuclei suspension buffer (10 mM Tris HCl, pH 7.4, 10 mM NaCl, 3 mM $MgCl_2$). Nuclei were filtered through a 30 µM strainer, then pelleted by centrifugation at 500 RCF for 5 minutes at 4°C. Supernatant was discarded and nuclei were resuspended in 500 µL of nuclei suspension buffer and pelleted again at 500 RCF at 4°C for 5 minutes. After discarding the supernatant the nuclei pellet was resuspended in 210 µL of nuclei suspension buffer. 2 aliquots of 100 µL nuclei suspension were snap-frozen in liquid nitrogen and stored in liquid nitrogen storage for sci RNA-Seq library preparation.

**Protein sample preparation.** 2.0 x $10^7$ of BAL cells, isolated AM, or isolated AT2 cells were lysed by suspension in 1 ml of lysis buffer containing protease and phosphatase inhibitors (RIPA Lysis Buffer System, Santa Cruz Biotechnology, Santa Cruz, CA) for 30 min on ice and centrifuged at 14,000 x *g* for 10 min, and the supernatant was collected. The concentrations of protein samples were measured using

Pierce™ BCA Protein Assay Kit (Thermo Scientific). The concentrations of RANKL in BAL cell lysate, AM lysate, or AT2 cell lysate were measured by ELISA and normalized to total protein concentrations.

**ELISA.** The levels of analytes in mouse BALF were measured by ELISA as follows: TRAP 5b, Mouse TRAP™ (TRAcP 5b) ELISA Kit (Immunodiagnostic Systems, United Kingdom); TNF-α, Mouse TNF alpha Uncoated ELISA Kit (Invitrogen, Waltham, MA); IL-1β , Mouse IL-1 beta Uncoated ELISA Kit (Invitrogen, Minneapolis, MN); IL-6 , Mouse IL-6 DuoSet ELISA Kit (R&D Systems Inc; IL-4, Mouse IL-4 DuoSet ELISA Kit (R&D Systems Inc); M-CSF, Mouse M-CSF DuoSet ELISA Kit (R&D Systems Inc); OPG, Mouse Osteoprotegerin/TNFRSF11B DuoSet ELISA Kit (R&D Systems Inc) according to the manufacturer's instructions. The levels of RANKL in mouse BAL fluid and and AT2 and BAL cell lysates were measured using the Mouse TRANCE/RANK L/TNFSF11 Quantikine ELISA Kit (R&D Systems Inc).

**Alveolar macrophage (AM) collection in mouse.** C57BL/6J mouse lungs were sacrificed at the indicated times post i.t. saline or silica challenge and BAL cells were collected. Cell pellets were resuspended in DMEM with 10% FCS and plated for isolation of AM by adherence to tissue culture plastic.

**Isolation of mouse and rat AT2 cells.** C57BL/6J mouse lungs were harvested on d14 post i.t. saline or silica challenge. Lungs were perfused with sterile normal saline via the pulmonary artery. The airway was cannulated via tracheostomy with an angiocatheter, and 2 ml of dispase (50 units/ml, Corning Inc., Corning, NY) was instilled, followed by 0.5 ml of 1% low-melt agarose (warmed to 45°C). Lungs were rapidly cooled on ice for 2 min, submerged in dispase for 45 min at room temperature, and transferred to a culture dish containing DNase I (0.1 mg/ml, Sigma-Aldrich). The parenchymal lung tissue was gently teased from the bronchi and homogenized. Cell suspensions were filtered, collected by centrifugation, and panned over prewashed 10 cm tissue culture plates coated with CD45 and CD32/16 antibodies (BD Biosciences, San Jose, CA). After incubation for 60 min at 37°C in a 5% $CO_2$ atmosphere to promote adherence of contaminating macrophages and fibroblasts, the AT2 cells were gently decanted from the plate, collected by centrifugation, and counted. Rat AT2 were isolated using published methods (*72*, *73*) with modifications. Animals were anesthetized with intraperitoneal sodium pentobarbital (75 mg/kg) and exsanguinated via the abdominal aorta. The right ventricle was cannulated and the lungs were perfused with heparin and flushed with 10ml Hepes buffered saline (HBS). The lungs were removed, lavaged with HBS + Ca, filled with 33 ml elastase solution in HPS + Ca (3U/ml) and incubated at 37 °C for 20 min. The lung parenchyma was cut into 1 mm pieces with scissors and incubated with 425 ul DNAase (Sigma #D4527; 17units/ml) in 10 ml of BSS-B at 37 °C with swirling. The suspension was filtered through 2 layers of gauze and 100 µm and 20 µm nylon filters to remove cell clumps and debris. The cells were centrifuged (8 min, 300×g, 4 °C) and the pellet was resuspended in buffer to remove red blood cells, and re-spun. The cell pellet was resuspended in DMEM-HEPES and panned over a rat IgG coated plate (Sigma # I4131) for 1 hour at 37°C in a 5 % CO2 atmosphere to remove macrophages. The supernatant was centrifuged (8 min, 300×g, 4 °C) to collect epithelial cells, and the supernatant was resuspended in 10ml of DMEM + 10% FBS, 1% penicillin/streptomycin and plated on collagen IV plates (Corning™ 354430).

**Lung digestion for flow cytometry.** The pulmonary vasculature was flushed with PBS (-) via right ventricular puncture. Lungs were removed and transferred to Iscove's

Modified Dulbecco's Medium (IMDM, Lonza, Malkersville, MD) containing 10% FCS, DNase I (0.1 mg/ml, Sigma-Aldrich), and 0.5 mg/ml of Liberase TL (Roche, Basel, Switzerland). Lungs were minced with scissors and incubated at 37°C for 45 minutes, after which time they were put on ice and pushed through a 70 µm strainer. Specimens were centrifuged at 1200 rpm at 4°C, resuspended in Red Blood Cell Lysing Buffer (Sigma-Aldrich) for 2 min, washed with media and filtered through 50 µm mesh. Cell viability was determined by trypan blue exclusion. Cells were plated into 96-well polystyrene round-bottomed culture plates at a density of $1x10^6$ cells/well in preparation for stimulation and staining.

**Flow cytometry.** For intracellular staining, the cells were treated with eBioscience ™ Cell stimulation cocktail (500X) consisting of phorbol 12-myristate 13-acetate (PMA) and ionomycin (Invitrogen, Walthman, MA), eBioscience ™ brefeldin A solution (1000X) (Invitrogen) and eBioscience ™ Monensin Solution (1000X) (Invitrogen) at 37°C for 2 hours. The cells with or without stimulation were stained with Live/Dead Aqua Dead Cell Stain Kit (Invitrogen), BUV395 Rat Anti-Mouse CD45 (BD Biosciences), BV605 Rat Anti-Mouse CD90.2 (BD Biosciences), APC anti-mouse CD3 (BioLegend, San Diego, CA), PE-Cy7 anti-mouse CD4 (BioLegend), PE-CF594 Rat Anti-Mouse CD8a (BD Biosciences), PerCP-Cyanine5.5 anti-mouse CD19 (BioLegend), FITC anti-mouse NK-1.1 ((BioLegend), PE anti-mouse CD254 (TRANCE, RANKL) (BioLegend), and BV421 Rat Anti-Mouse INF-γ (BD Biosciences) at 4°C, O/N. Four staining reactions per mouse were pooled before analyzing on an LSR Fortessa II cytometer (Becton, Dickinson and Company, Franklin Lakes, NJ). Data were analyzed with FlowJo software (TreeStar, Ashland, OR).

### Fig. S1, relates to Fig. 2

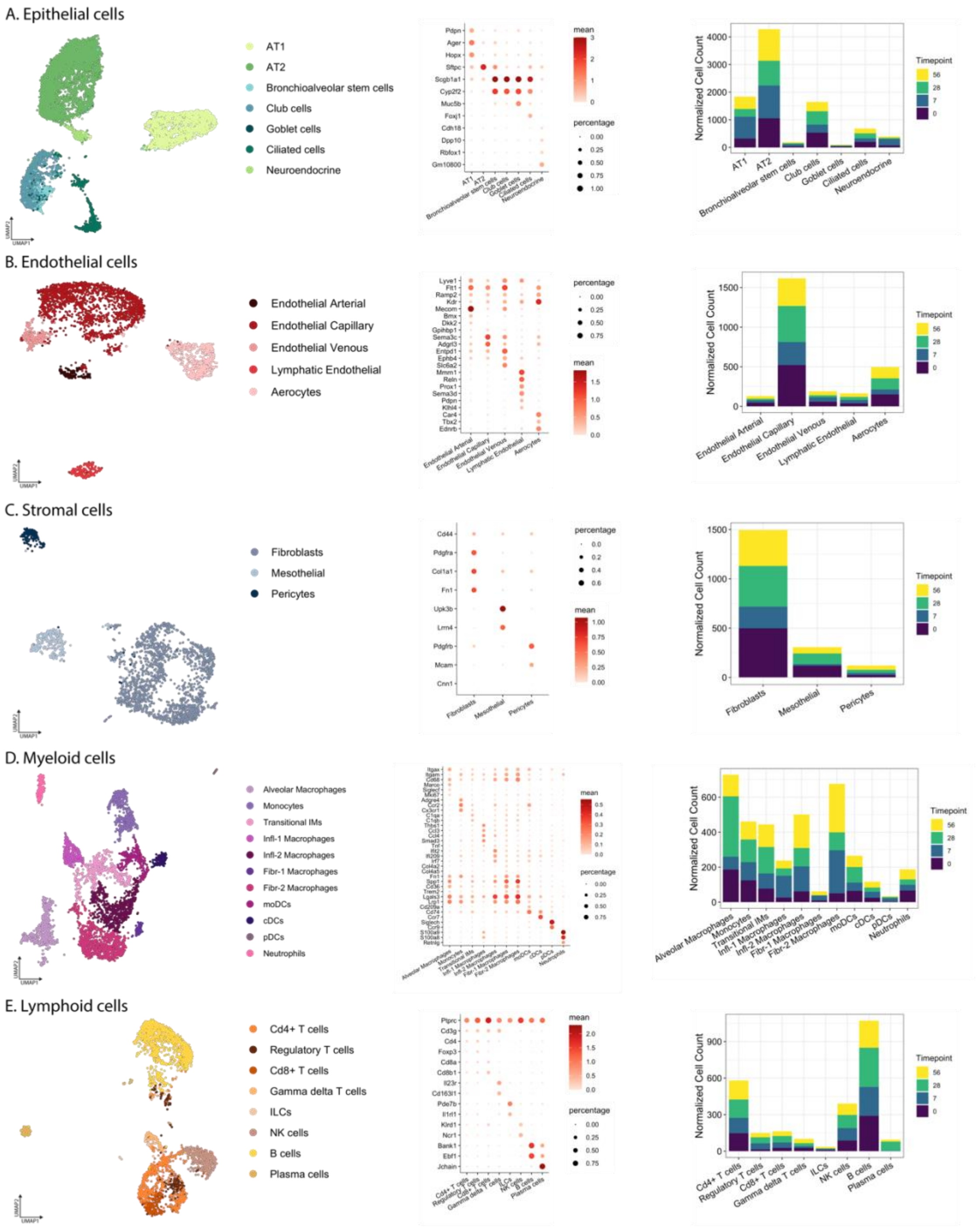


**Fig. S1. Fine annotation of single-nucleus RNA-sequencing data.** RNA from single nuclei were isolated and sequenced from Day 0 (pre-silica exposure) and Days 7, 28, and 56 (post-silica exposure). Cells were annotated based on clustering in reduced dimensions (left panels) and expression of marker genes (center panels) within major cell lineages in the murine lung: epithelial (A), endothelial (B), stromal (C), myeloid (D), and lymphoid (E). Total normalized number of each cell type are quantified and colored according to timepoint (right panels).

### Fig. S2, relates to Fig. 3

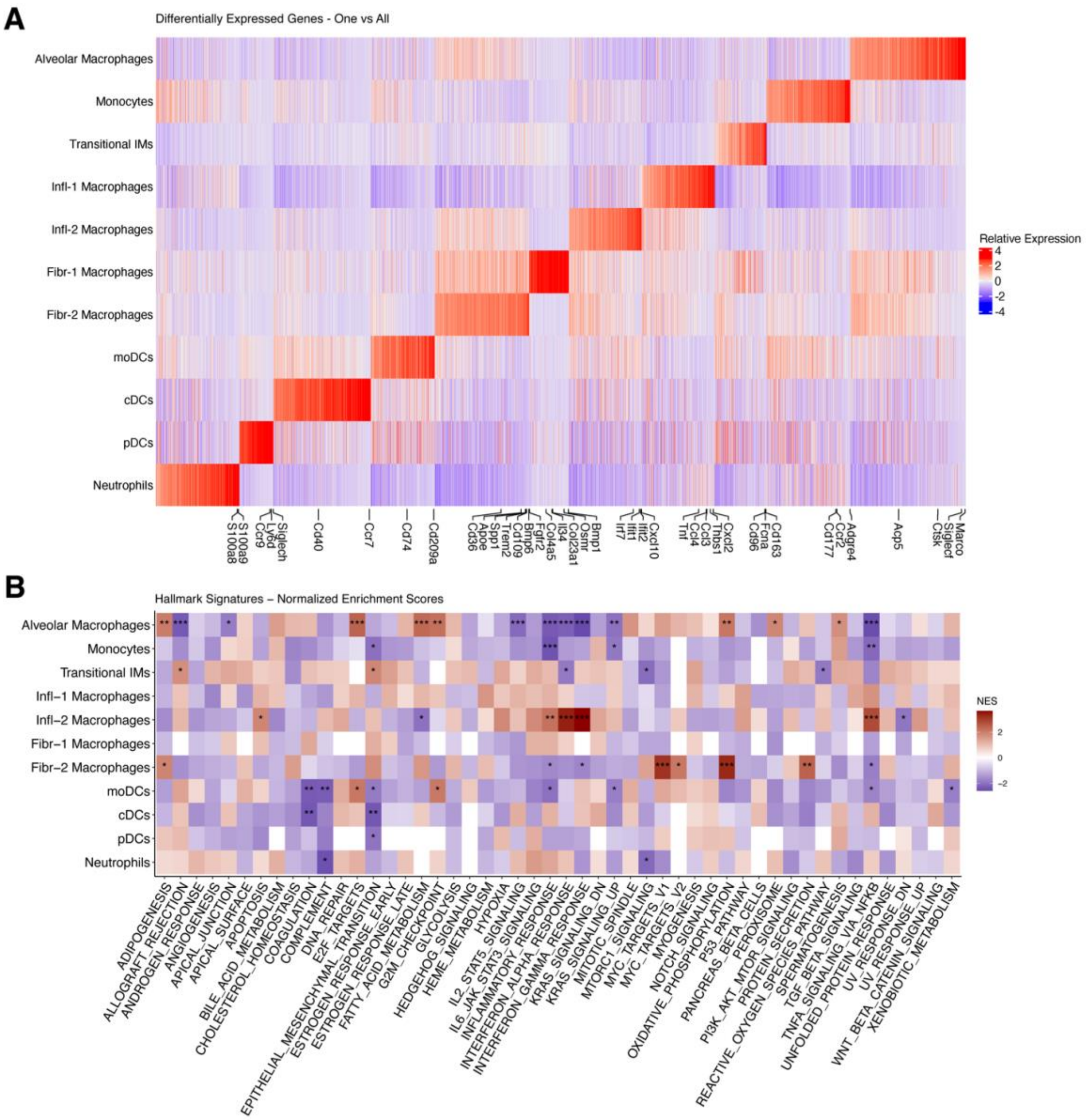


**Fig. S2. Analysis of gene expression and pathway activation differences to characterize myeloid cell states**. (A) For each cell state identified from clustering, we identified differentially expressed genes using linear mixed effect models and applying a one cell state vs all others approach. Genes were considered significant and retained for the analysis if the adjusted p-value <0.05 in any cell state. Genes were first grouped based on which cell state had the highest coefficient for the DEG and then plotted within these groups in descending order. (B) Normalized enrichment scores for Hallmark pathways demonstrate activation of various biological pathways in each cell state compared to other myeloid cell states. Normalized enrichment scores were calculated with fgsea using all coefficients from the DEG analysis. Significant NES are designated with asterisks (*p<0.05, **p<0.005, ***p<0.0005)

**Fig. S3**

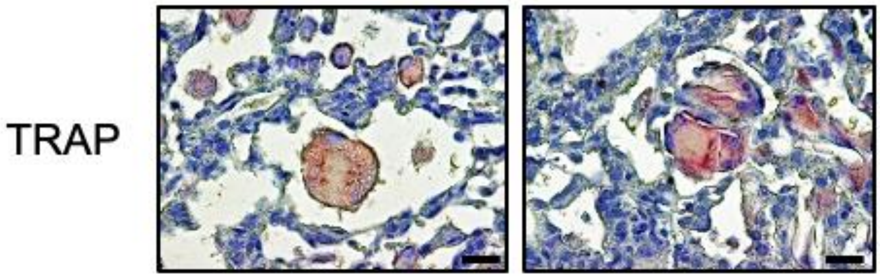


**Fig. S3. Silica-induced osteoclast-like cells persist up to 1 year post silica i.t. challenge.** Silica particles (5 mg) were administered i.t. into the lungs of C57BL/6J mice. Paraffin-embedded lung sections collected from mice 1 year post silica treatment were stained with TRAP (Scale bar, 20 μm).

**Fig. S4, relates to Fig. 4F**

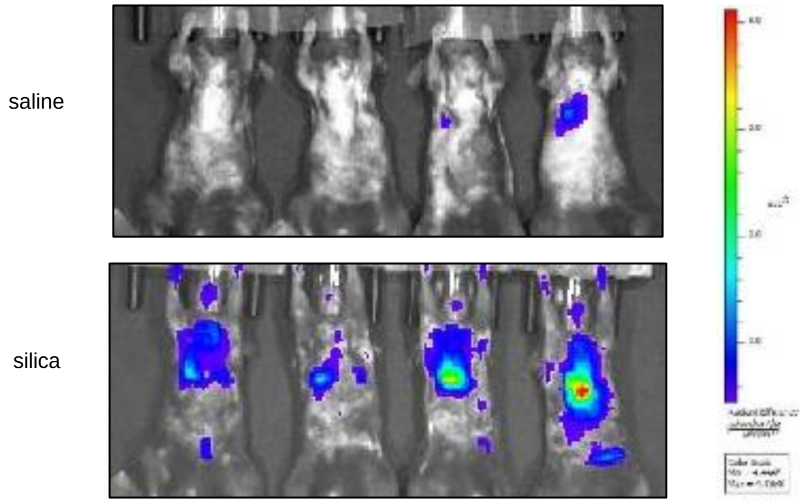


**Fig. S4. Intratracheal silica challenge induces CTSK enzyme activity in the lungs of mice**. Mice treated with saline or silica i.t. 6 days prior received the cleavage activated fluorescent cathepsin K substrate, Cat K 680 FAST by the i.t. route. Fluorescent images were acquired 18 h later by IVIS and (D) the fluorescent signal was quantified (N = 4 mice per group).

**Fig. S5, relates to Fig. 4**

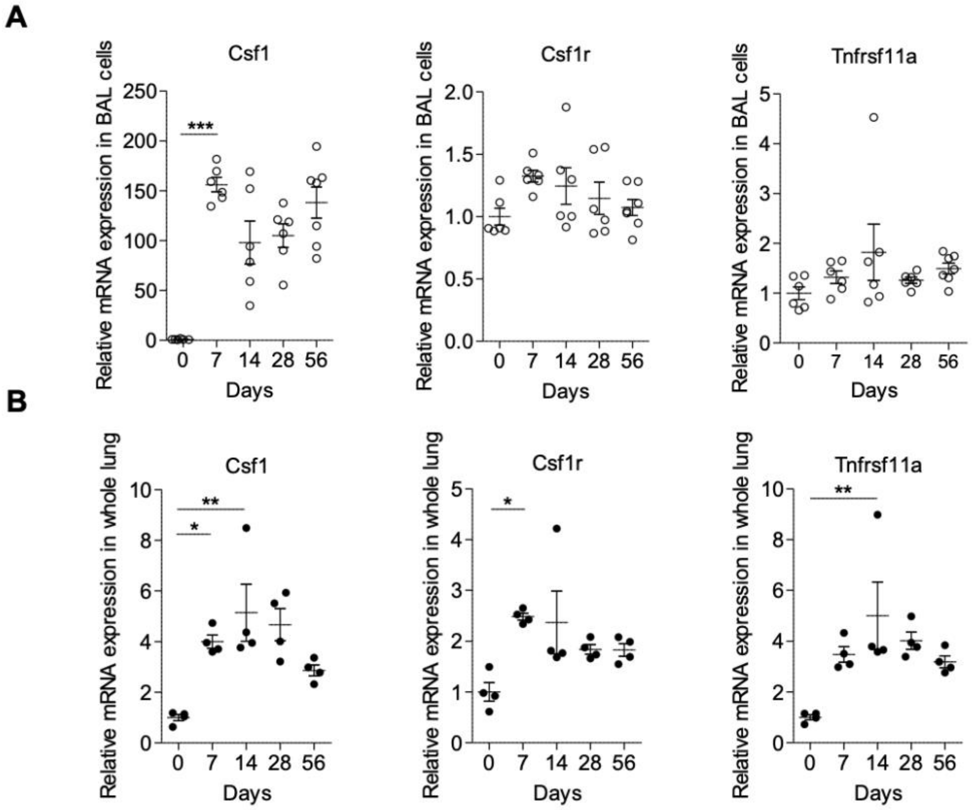


**Fig. S5. Osteoclast-related gene expression in BAL cells and lung tissue from silica exposed mice.** Silica particles (5 mg) were administered i.t. into the lungs of C57BL/6J mice. BAL cells (A, open circles) and whole lung tissue (B, black circles) were collected from mice at the indicated times in days (d) after administration, and osteoclast-related gene expression was assessed by rtPCR for csf1 (M-CSF), csf1r (M-CSF receptor), and Tnfrsf11a (RANK) (N = 6-7 mice per group). *$P$<0.05, **$P$<0.01, ***$P$<0.001

**Fig. S6**

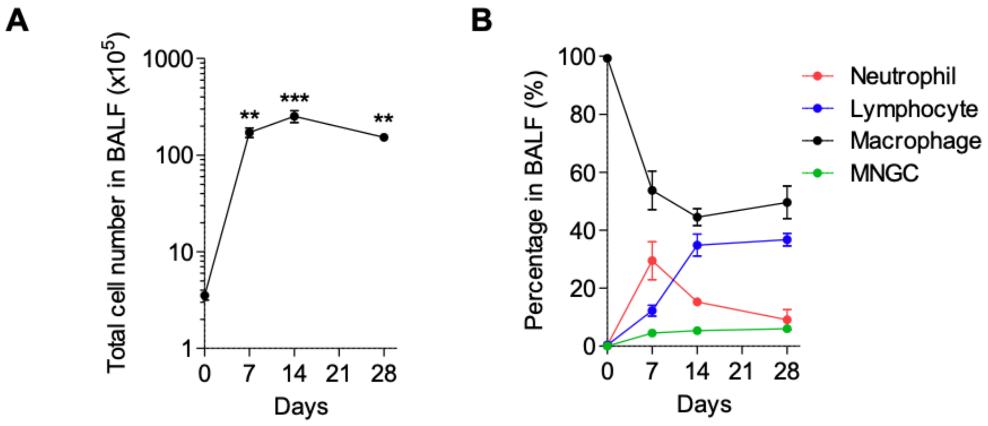


**Fig. S6. BAL cell numbers and differential in i.t. silica challenged mice.** Silica particles (5 mg) were administered i.t. into the lungs of C57BL/6J mice. (A) The number BAL cells at indicated times in days (d) after silica challenge are shown (N = 4 mice per group). (B) The BAL cellular composition at indicated times in days (d) after silica challenge are shown (N = 4 mice per group). *$P$<0.05, **$P$<0.01, ***$P$<0.001

**Fig. S7, relates to Fig. 4I and 4J**

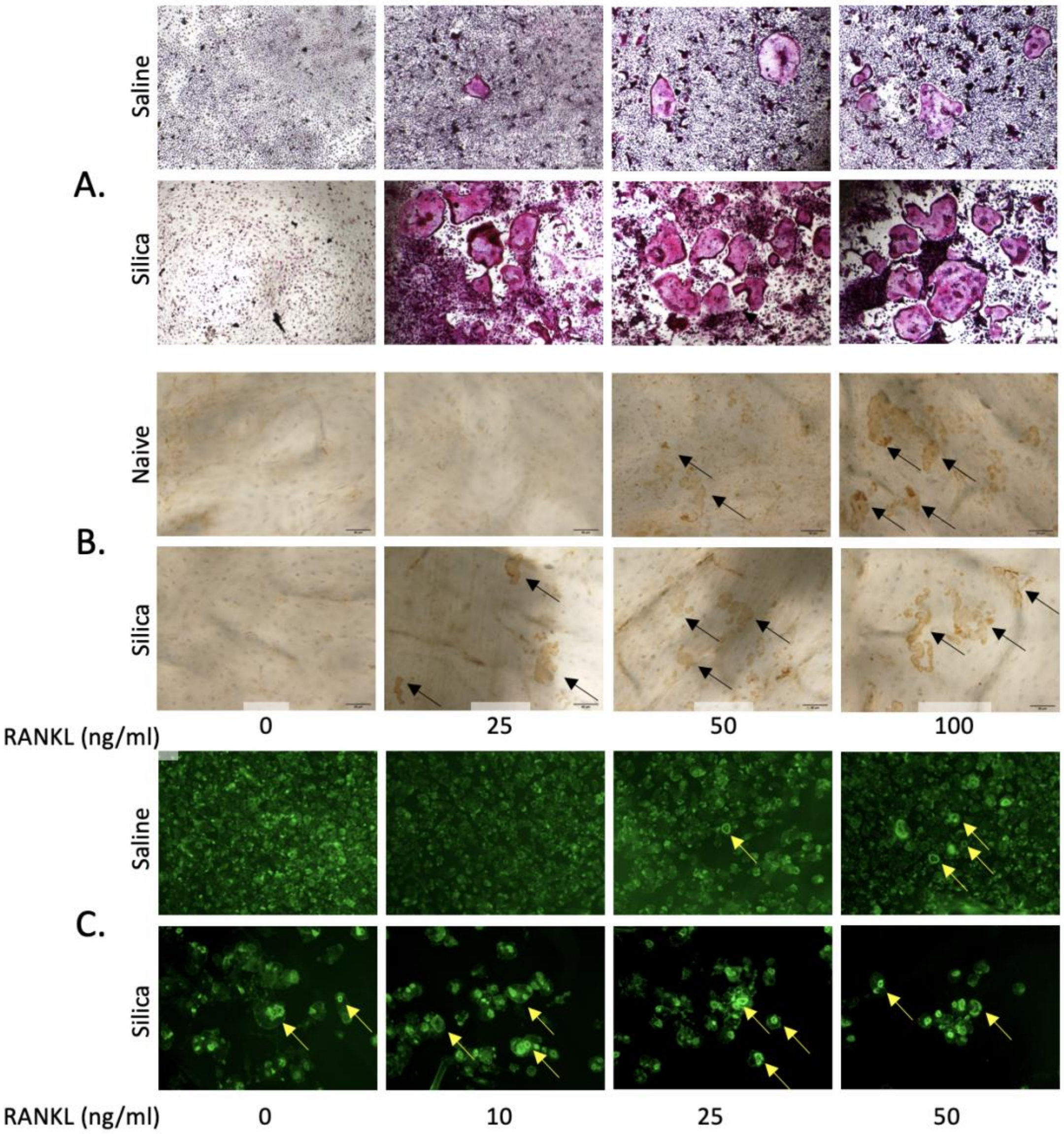


**Fig. S7. Intratracheal silica challenge enhances osteoclast formation, bone pitting and actin ring assembly is cultured BAL cells.** BAL cells isolated from C57BL/6J mice at day 14 post intratracheal challenge with saline or silica were plated on plastic plates or bovine bone slices with M-CSF and RANKL at indicated concentrations. At day 6, the cells cultured on the plastic plates were stained for TRAP activity (Panel A). For BAL cells plated on bone slices, resorbed bone area was visualized by peroxidase-conjugated wheat germ agglutinin/horse radish peroxidase staining (black arrows, Panel B). Actin rings were visualized by phalloidin staining (yellow arrows, Panel C).

**Fig. S8, related to Fig. 4I and 4J**

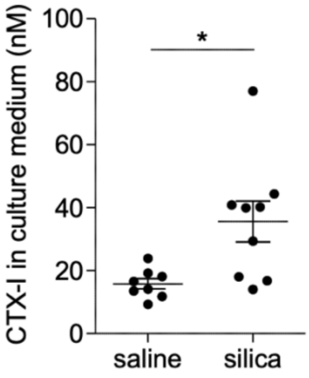


**Fig. S8. Intratracheal silica challenge enhances bone matrix degradation by BAL cells cultured on bone slices.**

BAL cells isolated from C57BL/6J mice at day 14 post intratracheal challenge with saline (saline) or silica were plated on bovine bone slices as above. The levels of CTX-I in bone culture medium were measured by ELISA (N=8-9 wells per group). *$P<0.05$, **$P<0.01$, ***$P<0.001$

**Fig. S9, related to Fig. 5**

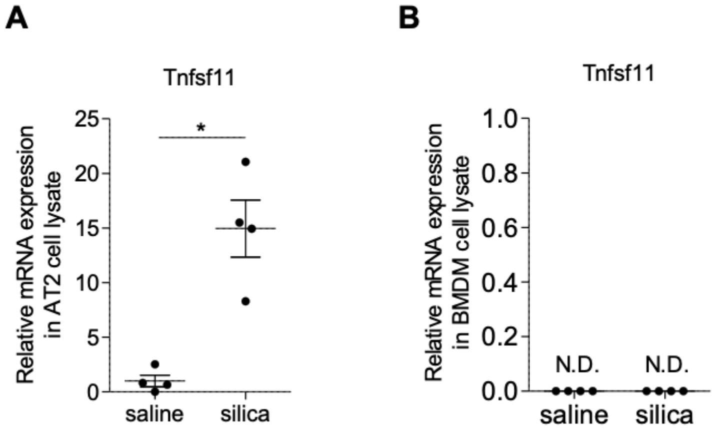


**Fig. S9. Silica particles induce RANKL expression in primary rat AT2 cells.**
(A) Isolated rat AT2 cells were incubated in DMEM with 10% (v/v) FCS in the presence or absence of 10 µg/ml of silica particles for 3 days. Relative gene expression of Tnfsf11 (RANKL) in AT2 cell lysate was evaluated by rtPCR (N = 4 wells per group). (B) Isolated mouse BMMs were incubated with DMEM in 10% (v/v) FCS and M-CSF (5 ng/ml) in the presence or absence of 10 µg/ml of silica particles for 3 days. Relative gene expression of Tnfsf11 (RANKL) in BMMs was evaluated by rtPCR (N = 4 wells per group) (N.D.; not detected). ***P*<0.01.

### Table S1

| Gene | Primer | |
|---|---|---|
| | Forward | Reverse |
| Acp5 | 5’-GCCACAGTTATGTTTGTACGTG-3 | 5’-ACAGATTGCATACTCTAAGATCTCC-3′ |
| Acta2 | 5′-CTGTTATAGGTGGTTTCGTGG A-3′ | 5′-GAGCTACGAACTGCCTGAC-3′ |
| Actb | 5′-ACCTTCTACAATGAGCTGCG-3′ | 5′-CTGGATGGCTACGTACATGG-3′ |
| Atp6v0d2 | 5’-GCCAAATGAGTTCAGAGTGATG-3’ | 5’-AGTCTTACCTTGAGGCATTCTAC-3’ |
| Col1α1 | 5’-CATTGTGTATGCAGCTGACTTC-3’ | 5’-CGCAAAGAGTCTACATGTCTAGG-3’ |
| Col3α | 5‘-TCTCTAGACTCATAGGACTGACC-3’ | 5’-TTCTTCTCACCCTTCTTCATCC-3’ |
| Csf1 | 5’-GGAAGATGGTAGGAGAGGGTA-3’ | 3’-AGGATGAGGACAGACAGGT-5’ |
| Csf1r | 5′-AGGTGTAGCTATTGCCTTCG-3′ | 5′-TGTATGTCTGTCATGTCTCTGC-3′ |
| Ctsk | 5’-ATCTCTCTGTACCCTCTGCAT-3′ | 5’-GACTCTGAAGATGCTTACCCA-3′ |
| Fn1 | 5’-TTGTTCGTAGACACTGGAGAC-3’ | 5’-GAGCTATCCATTTCACCTTCAGA-3’ |
| Itgb3 | 5’-ACAGTCATCCTCGTTCTTGTAG -3′ | 5’-GAACGCTCCATGAAGAAAACAC-3′ |
| Mmp9 | 5’-GTGGGAGGTATAGTGGGACA-3′ | 5’-GACATAGACGGCATCCAGTATC-3′ |
| Tgfb1 | 5’-CCGAATGTCTGACGTATTGAAGA-3’ | 5’-GCGGACTACTATGCTAAAGAGG-3’ |
| Tnfsf11 | 5’-AGTGCTGTCTTCTGATATTCTGT -3′ | 5′-TCCCGCTCCATGTTCCT-3′ |
| Tnfrsf11a | 5’-CACTGTCGGAGGTAGGAGT-3’ | 5’-CAGGAGAGGCATTATGAGCAT-3’ |
| Tnfrsf11b | 5’-ATGCAACACATGACAACGTG-3′ | 5′-TGGTATAATCTTGGTAGGAACAGC-3′ |

**Table S1. Primers for quantitative RT-qPCR**

Acp5, acid phosphatase 5; Acta2, actin α-2 smooth muscle; Actb, β-actin; Atp6v0d2, ATPase $H^+$ transporting v0 subunit d2; Col1α1, collagen 1α-1; Col3α1, collagen 3α-1; Csf1, colony stimulating factor 1; Csf1r, Csf1 receptor; Ctsk, cathepsin K; Fn1, fibronectin 1; Itgb3, Integrin β3; Mmp9, matrix metallopeptidase 9; Tgfb1, transforming growth factor-β1; Tnfsf11, tumor necrosis factor receptor superfamily member 11; Tnfrsf11a , tumor necrosis factor receptor superfamily member 11a; Tnfrsf11b, tumor necrosis factor receptor superfamily member 11B
